# Human hippocampal ripples prioritise model-based learning

**DOI:** 10.1101/2025.07.31.667862

**Authors:** Xiaoyu Zhou, Xiongfei Wang, Xiangyu Hu, Haiteng Wang, Jinbo Zhang, Qianqian Yu, Jiahua Xu, Zhibing Xiao, Li He, Yunzhe Liu

## Abstract

Humans often learn optimally, inferring the value of options they have never directly experienced by leveraging internal models of the world, a process known as model-based learning. Yet how the brain decides which unexperienced options to update first remains unclear. Here we recorded intracranial EEG from 34 epilepsy patients performing a reinforcement-learning task that required using task structure to infer the value of unvisited (non-local) paths. After each reward, brief hippocampal “ripple” events signalled which indirect experience was most valuable, thereby encoding that path’s priority. Longer ripples carried the strongest priority signals, allowing the brain to update high-value, unvisited options first and thereby optimise learning. Ripple events also coincided with selective cortical reactivation of these high-priority paths, consistent with prioritised replay. Crucially, when hippocampal ripples were precisely synchronised with activity in the lateral frontopolar cortex (LFPC), individuals showed greater model-based learning: stronger ripple–LFPC coupling predicted more effective use of task structure and more accurate learning of non-local values. Our findings establish hippocampal ripple-centred prefrontal coordination as a fundamental mechanism that prioritises valuable experiences for model-based learning, explaining how the human brain learns so efficiently.

## Introduction

When we encounter new experiences, we must learn optimally: drawing on what we already know to infer what might happen next so that a single encounter can guide many future choices. Imagine you discover a vein of gold in a remote valley. You soon favour not only the route you just walked but also alternative paths that converge on that valley, despite never having taken them.

In reinforcement learning (RL), this capacity is formalised as model-based learning (Daw et al., 2005; Doll et al., 2012), in which indirect (“non-local”) outcomes are inferred from direct (“local”) experiences and used to guide future choices. In essence, the brain is able to learn value of unvisited options – a non-local value update – using its internal model of the environment.

Rodent studies have highlighted the hippocampus as central to model-based learning (Miller et al., 2017). During rest or pauses in navigation, hippocampal neurons spontaneously replay sequences of both visited and unvisited locations (Diba & Buzsáki, 2007; Gupta et al., 2010; Karlsson & Frank, 2009; A. K. Lee & Wilson, 2002; Ólafsdóttir et al., 2015, 2018; Pfeiffer & Foster, 2013; Widloski & Foster, 2022). In humans, non-invasive recordings indicate that such replay drives value propagation to non-local paths (Y. Liu, Mattar, et al., 2021). Crucially, replay is not random: it is biased towards trajectories with high “priority”, so that the most useful indirect experiences are updated first, an optimisation predicted by RL theory (Agrawal et al., 2022; Mattar & Daw, 2018). Yet how the brain implements this prioritisation to optimise learning, remains unknown.

Hippocampal ripples offer a promising cellular mechanism (Krausz et al., 2023). Ripples are brief bursts of high-frequency oscillations reflecting coordinated neuronal firing. In rodents, ripples coincide with replay (Bush et al., 2022; Buzsáki, 2015), coordinate with midbrain reward-responsive neurons (Gomperts et al., 2015), and support reward learning in both physical (Foster & Wilson, 2006; Ólafsdóttir et al., 2015) and abstract spaces (Barron et al., 2020). Ripples also contain representations of the environment (Widloski & Foster, 2024; Wu & Foster, 2014), and of previously rewarded or unvisited locations (Gillespie et al., 2021), suggesting they could integrate reward signals with the world model to guide non-local value updates.

Model-based learning also depends on hippocampal–prefrontal interactions. Human fMRI work on decision-making implicates the medial prefrontal cortex (PFC), lateral orbitofrontal cortex (LOFC), and particularly the lateral frontopolar cortex (LFPC) in representing task structure, assigning credit, and evaluating alternative options (Boorman et al., 2009, 2011; Daw et al., 2011; Doll et al., 2015; S. W. Lee et al., 2014; Moneta et al., 2023; Schuck et al., 2016). Interactions between these prefrontal regions and the hippocampus are also frequently observed (Boorman et al., 2016; Garvert et al., 2023; Muhle-Karbe et al., 2023; Schuck & Niv, 2019; Witkowski et al., 2024). However, these neuroimaging studies offer indirect measures of neuronal activity, and cannot directly test their functional circuitry with hippocampus. Rodent work shows that hippocampal ripples synchronise with prefrontal regions (Berners-Lee et al., 2021; Nitzan et al., 2022), and can modulate prefrontal representations on-task (Jensen et al., 2024). Yet the precise role of ripple-aligned prefrontal activity in model-based learning remains largely unknown.

Here we addressed these questions by combining a validated model-based RL task (Y. Liu, Mattar, et al., 2021) with intracranial electrophysiology (iEEG) in 34 epilepsy patients. This approach allowed us to directly record neuronal activity simultaneously from hippocampus (157 contacts), and cortical areas (2,536 contacts) during learning. We identify a priority-based learning mechanism, orchestrated by hippocampal ripples and enacted through prefrontal cortex, that explains how the human brain learns so efficiently.

## Results

### Experimental design

We recruited 34 individuals (20 males and 14 females, ages 26.3 ± 1.7 years, mean ± SEM; **Extended Data Table 1**) with pharmacologically resistant epilepsy, who underwent iEEG recording for clinical evaluation.

They performed a three-armed bandit-style RL task requiring the use of an internal model of the task structure (**Fig. 1a**) (Y. Liu, Mattar, et al., 2021). Each trial began with one of three possible “arm” cues, which branched into two distinct paths leading to one of two possible end states (**Fig. 1b**). Subjects were instructed to maximise reward and, to do so, needed to leverage the task structure (i.e. knowledge of which paths lead to which end states). This design dissociates local learning – updating the value of the chosen path – from non-local learning, which is the propagation of value to the other paths that share the same end state.

**Fig. 1.**
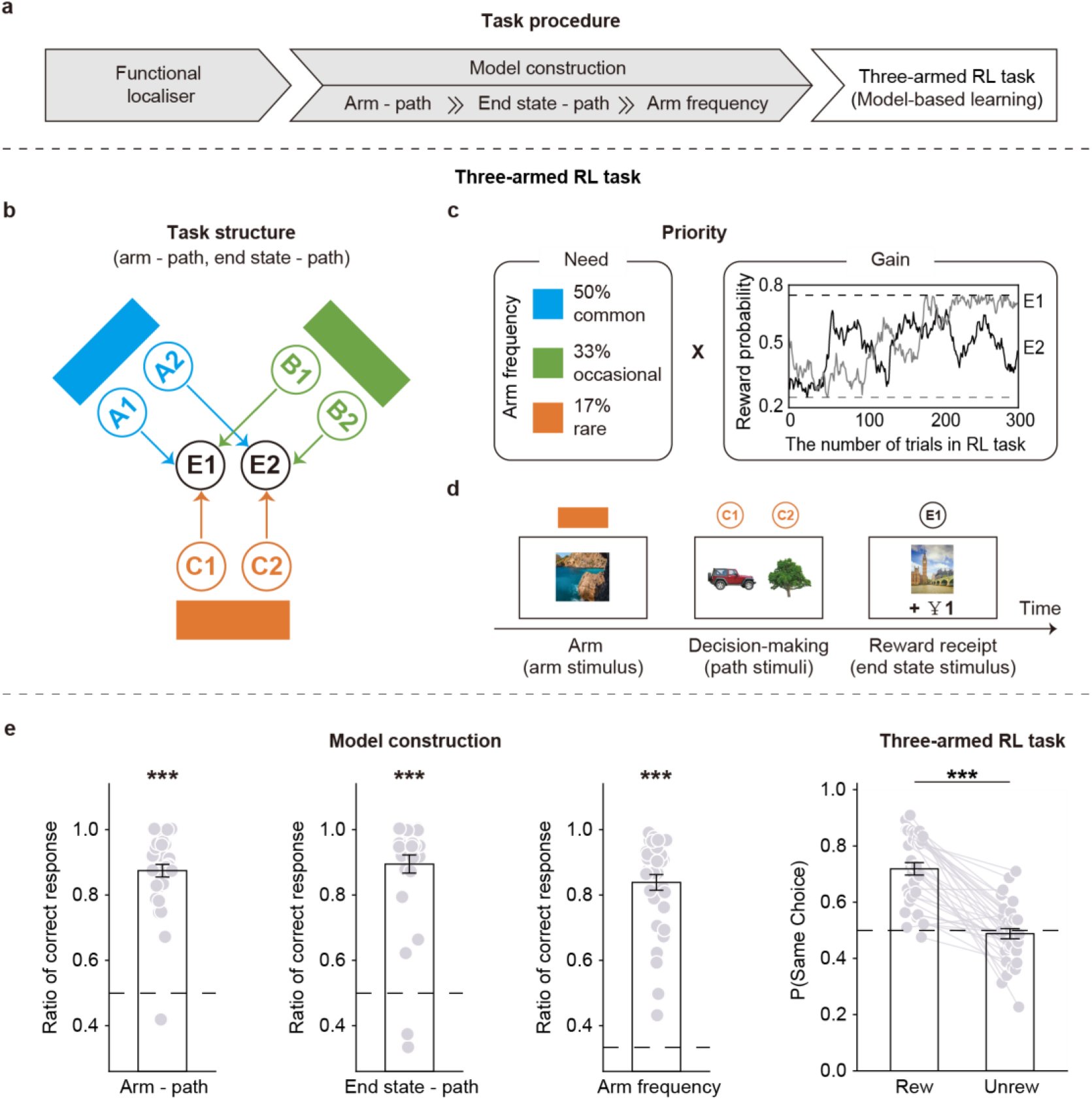
Experimental design and behavioural performance. **a**, Experimental procedure adapted from a previous study (Y. Liu, Mattar, et al., 2021). Subjects first completed the functional localiser to identify neural signatures of visual stimuli (for later use in the RL task; see Methods). They then learned task structure about, arm-path mappings, end-state-path mappings, and arm frequency. This learned task-related information is an essential prerequisite for non-local learning and need manipulation. Finally, they completed the RL task, which required them to utilise the acquired task structure to maximise reward. **b**, Task structure in the three-armed RL task. Each arm consistently links to two paths, and each end state consistently links to three paths, allowing investigation of non-local learning, i.e. experience in one visited path influences the other two unseen, non-local paths sharing the same end state. **c**, Manipulation of priority. Priority measures the utility of value propagation in each non-local path, defined as the product of need and gain, according to RL theory (Agrawal et al., 2022; Mattar & Daw, 2018). The need refers to how often each arm appears (fixed arm frequencies, as learned in the model construction task). The gain varies trial by trial, due to the consistent shifts in reward probabilities of end states E1 and E2 throughout the RL task. These probabilities follow independent Gaussian random walks between 25% and 75%. **d**, Illustration of a core trial in the RL task. First, the Arm screen – one of the three arms is displayed according to its predetermined probability. Next, the Decision-making screen – two arm-specific paths are shown and the subject selects one. Lastly, the Reward receipt screen – the subject sees whether they received a reward (¥1) or not (¥0). **e**, Task performance. Subjects showed good knowledge of the task structure (arm–path and end-state–path mappings) and arm frequencies. In the RL task, subjects preferred the path that led to the previously rewarded end state. “Rew/Unrew” denotes last-trial reward status. P(Same Choice) is the proportion of trials in which participants selected a path leading to the same end state as on the preceding trial. The dashed line indicates chance level. Error bars show SEM across subjects. Each dot represents one subject. Each grey line connects data from the same subject. *** *p* < 0.001.

Importantly, non-local learning was hypothesised to be modulated by the priority of accessing states offline (Agrawal et al., 2022; Antonov et al., 2022; Antonov & Dayan, 2025; Mattar & Daw, 2018). According to RL theory (Agrawal et al., 2022; Mattar & Daw, 2018), this priority was defined as the product of “*need*” and “*gain*” (**Fig. 1c**). The “*need*” reflected how frequently each arm was presented (rare-17%, occasional-33%, common-50%), while the “*gain*” depended on prediction errors (PE) and “value of information” that increased when an arm remained unvisited for multiple trials (see Methods for details). Note the reward probabilities of both end states changed slowly and independently (random walks between 25% and 75%), thereby enabling consistent learning. Together, these features allowed us to investigate how learning extends beyond direct experience to nonlocal paths, and how it is prioritised in the brain.

The full task protocol consisted of a functional localiser task, a model construction task, and the main three-armed RL task (**Fig. 1a**). The functional localiser task allowed us to train stimulus-specific neural decoders to detect subsequent task-related reactivations. In this task, six distinct visual stimuli—later serving to index “paths” in the RL task—were presented in random order. Subjects were asked to indicate whether a displayed word matched the preceding stimulus. The mean accuracy was 0.984 ± 0.002 (t(26) = 197.54, *p* = 7.8 × 10^-43^, one-sample *t*-test against chance), suggesting that subjects were highly attentive. Seven subjects did not participate this localiser and were thus excluded from reactivation analyses, although they remained in the broader behavioural and neural assessments.

The model construction task served two purposes. First, it ensured that subjects had a comprehensive understanding of the task structure underpinning the three-armed RL task. This understanding is a prerequisite of non-local learning. Subjects first learned deterministic mappings between each arm and its two specific paths (arm-path mapping), then learned how each end state was linked to three such paths (end state-path mapping). These mapping tasks were repeated until subjects met predetermined criteria (see Methods for details). Second, the task established each arm’s fixed appearance probability (arm frequency), which later defined the “need” manipulation in the main RL task. In the final test blocks (**Fig. 1e**), the mean accuracies were 0.874 ± 0.019 in the arm-path mapping (t(33) = 19.44, *p* = 1.2 × 10^-19^, one-sample *t*-test against chance), 0.894 ± 0.028 in the end state-path mapping (t(33) = 14.19, *p* = 1.3 × 10^-15^), and 0.838 ± 0.024 in the arm frequency (t(33) = 21.06, *p* = 1.0 × 10^-20^), showing that subjects successfully acquired the relevant knowledge of task structure and need.

After completing these tasks, subjects performed the main three-armed RL task over 300 trials, during which prioritised non-local learning was investigated. Each trial began with a single arm, which was chosen pseudo-randomly according to its assigned arm frequency. Two path stimuli from that arm then appeared, and participants selected one path at their own pace to observe the corresponding end state. They then pressed a key to proceed to the reward receipt screen, which displayed either ¥0 or ¥1 (**Fig. 1d**). As a sanity check, a mixed-effects logistic regression showed that participants were significantly more likely to select the same end state in which they had just been rewarded than if they had not been rewarded (*p* = 4.0 × 10^-10^; **Fig. 1e**), indicating effective learning from reward feedback.

### Behavioural evidence of prioritised non-local learning

In our task, local vs. non-local learning make different behavioural predictions. If learning is purely local, a reward should only influence future choices when the same arm appears again (since only the experienced path’s value is updated). In contrast, model-based (non-local) learning predicts that a reward will also affect choices even when a different arm is presented next, as long as that arm leads to the same end state (because the value of the unexperienced path to that end state should be inferred).

Although subjects successfully learned the task structure, their knowledge was not fully accurate (**Fig. 1e**). Moreover, humans might not perfectly utilise the task structure in every trial – they might sometimes generalise a reward inappropriately to unseen paths associated with the other end state (**Fig. 2a**, dashed purple arrows). Such “confused” updating can make non-local learning appear less effective than local learning. This would manifest as an “efficiency gap”: choices would be influenced more strongly by recent rewards when the arm repeats (local) than when it switches (non-local).

**Fig. 2.**
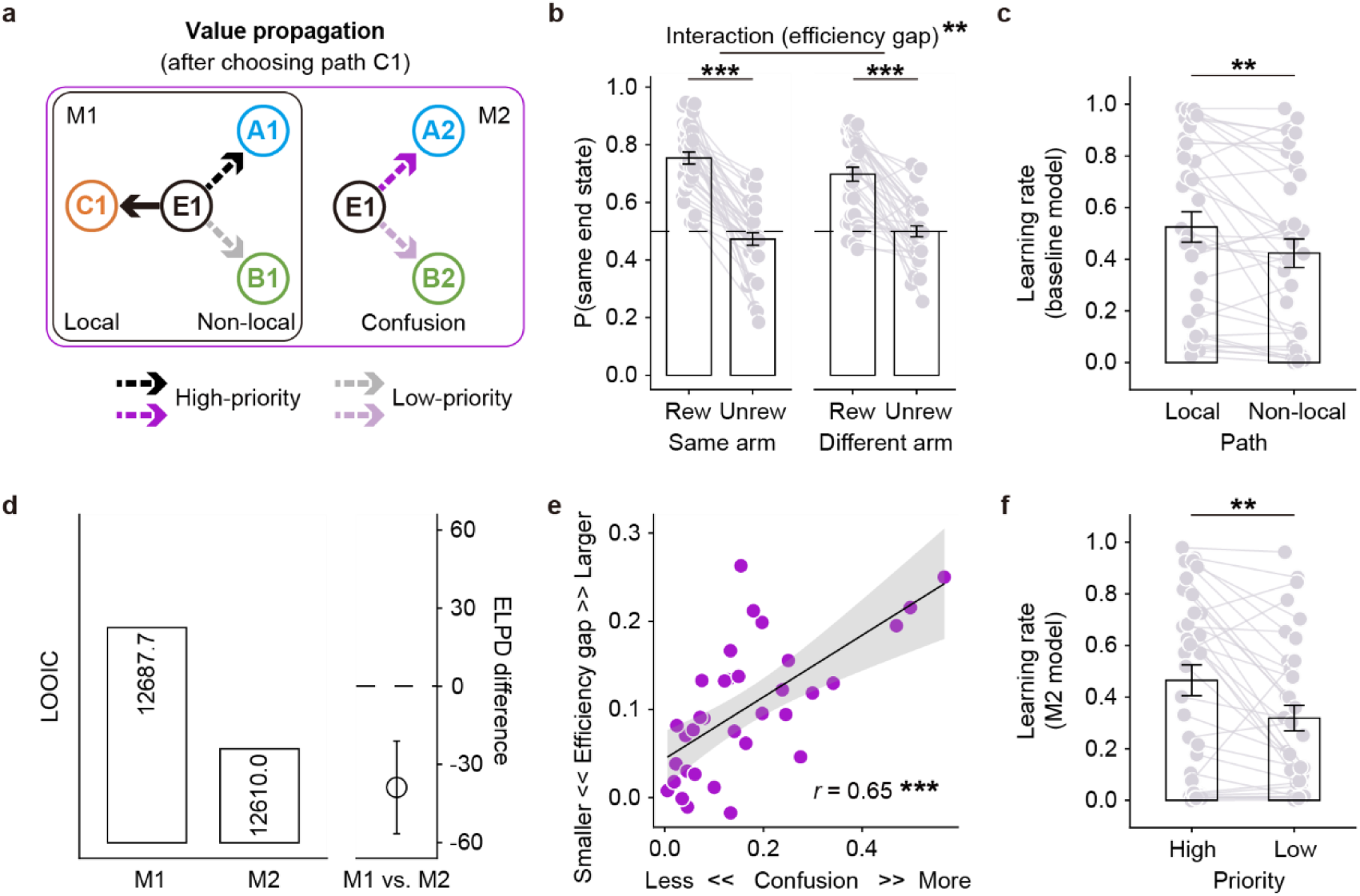
Behavioural evidence of prioritised non-local learning. **a**, Illustration of prioritised non-local learning with imperfect structure utilisation. Both models M1 and M2 assume that learning occurs in both local and non-local paths, and that learning in non-local paths can differ based on their priority. For example, receiving a reward at E1 via path C1 results in value propagation for both C1 (local learning, solid arrow) and its associated upstream paths A1 and B1 (non-local learning, dashed black/grey arrows). M2 additionally assumes that if subjects do not perfectly utilise knowledge of the task structure, they might also incorrectly update unseen paths to the other end state (e.g. A2 and B2, dashed light/dark purple arrows), in proportion to a confusion parameter, χ. **b**, Model-agnostic evidence of non-local learning. Subjects’ preference for a path was higher if the end state associated with it was rewarded in the previous trial, even when the current trial began with a different arm (non-local learning). However, this reward effect was stronger when the same arm repeated (local learning) suggesting that subjects learn less effectively on non-local paths than local ones. This interaction was referred to as ‘efficiency gap’. ‘Rew/Unrew’ denotes last-trial reward status. P(Same Choice) is the proportion of trials in which participants selected a path leading to the same end state as on the preceding trial. **c**, RL modelling results. The learning rates for non-local paths were significantly lower than for local paths, consistent with the model-agnostic evidence in panel **b**. **d**, Model comparison. The augmented model M2 (incorporating imperfect structure utilisation, confusion parameter, χ) provided a better fit to behaviour than the original priority model M1. **e**, The confusion parameter is linked to the efficiency gap. Specifically, the confusion parameter, χ, estimated from M2 correlated positively with the logistic regression interaction term measuring the efficiency gap. **f**, In M2, high-priority non-local paths had higher learning rates than low-priority non-local paths, indicating prioritised non-local learning. Each grey line connects data from the same subject. Error bars show SEM. ** *p* < 0.01, *** *p* < 0.001.

We analysed choices as a function of the previous trial’s outcome (rewarded vs. not rewarded), whether the arm changed between trials (same vs. different arm), and their interaction, using a mixed-effects logistic regression. In keeping with prior findings (Y. Liu, Mattar, et al., 2021), subjects were more likely to select a path leading to a previously rewarded end state whether or not the arm changed, indicating that both local and non-local learning operated (reward effect at the “same arm” level: *p* = 1.8 × 10^-19^, reward effect at the “different arm” level: *p* = 5.1 × 10^-12^). However, this reward effect was weaker when the arm changed than when it was repeated (*p* = 1.2 × 10^-3^, for the interaction between last-trial outcome and arm transition; **Fig. 2b**), indicating an efficiency gap between local and non-local learning.

The analysis above considered only the immediate reward, overlooking longer time horizon of reward history. To capture this, we built a computational model that jointly captured local and non-local learning across all trials within a single RL framework (see (see Methods for details). In this baseline model, values in both paths were updated from received rewards, with separate learning-rate parameters for local (*α*_*l*_) and non-local (*α*_*nl*_) paths. As in the regression analysis, non-local learning rates were significantly lower than local learning rates (*α*_*l*_: 0.525 ± 0.059, *α*_*nl*_: 0.423 ± 0.055; paired *t*-test, t(33) = 3.19, *p* = 3.1 × 10^-3^; **Fig. 2c**). Together, the regression and computational results show that subjects did learn in a non-local manner, but less effectively than from direct experience.

Next, we asked whether non-local learning varied with the priority, according to normative RL theory (Agrawal et al., 2022; Mattar & Daw, 2018). We expanded the baseline model by replacing non-local learning rate (*α*_*nl*_) with two different parameters (*α*_*high*_ for high-priority path, *α*_*low*_ for low-priority path; **Fig. 2a**, dashed black/grey arrows), referred to as M1. Priority was defined as the product of need and gain (see Methods for details).

A plausible reason for reduced efficiency is “confused” updating driven by imperfect structure utilisation (Antonov et al., 2022; Costa et al., 2023; Vikbladh et al., 2019). To test this, we further extended M1 to include imperfect structure utilisation, producing M2 (Costa et al., 2023). In both M1 and M2, *α*_*high*_ and *α*_*low*_ varied with computed priority (need × gain). M2 additionally assumed that imperfect knowledge could lead subjects to retrieve incorrect paths leading to the other end state (**Fig. 2a**, dashed purple arrows), updating those paths in proportion to a “confusion” parameter *χ*. When *χ* = 0, M2 reduces to M1.

Model comparison showed that M2 fit the behaviour better (expected log pointwise predictive density, ELPD difference: −38.8, SE: 17.8; **Fig. 2d**; see also **Extended Data Fig. 1** for parameter convergence). Moreover, *χ* correlated positively with the logistic-regression estimated “efficiency gap” (r = 0.65, *p* = 2.9 × 10^-5^; **Fig. 2e**), indicating that poorer structure utilisation produced a larger gap. Finally, in M2 the learning rate for high-priority non-local paths was significantly higher than for low-priority paths (*α*_*high*_: 0.465 ± 0.060, *α*_*low*_: 0.319 ± 0.050, paired *t*-test, t(33) = 3.58, *p* = 1.1 × 10^-3^; **Fig. 2f**), showing that participants prioritised non-local value updates according to our priority metric, effectively learning more from rewarding experiences that the task model deemed more important.

### Hippocampus and prefrontal cortex are involved in the non-local learning

Having established behavioural evidence for non-local learning and its prioritisation, we next examined the underlying neural mechanisms. Three participants lacked hippocampal electrodes and were excluded, leaving 31 patients with depth electrodes in both the hippocampus (157 contacts) and widespread cortex (2,536 contacts, including 839 prefrontal contacts; **Fig. 3a, b**; see also **Extended Data Fig. 2a** and **Extended Data Table 2**). Throughout we analysed high-frequency broadband (HFB; 60–160 Hz) power (**Extended Data Fig. 2b, c**), a reliable proxy for local population spiking (Mukamel et al., 2005; Parvizi & Kastner, 2018).

**Fig. 3.**
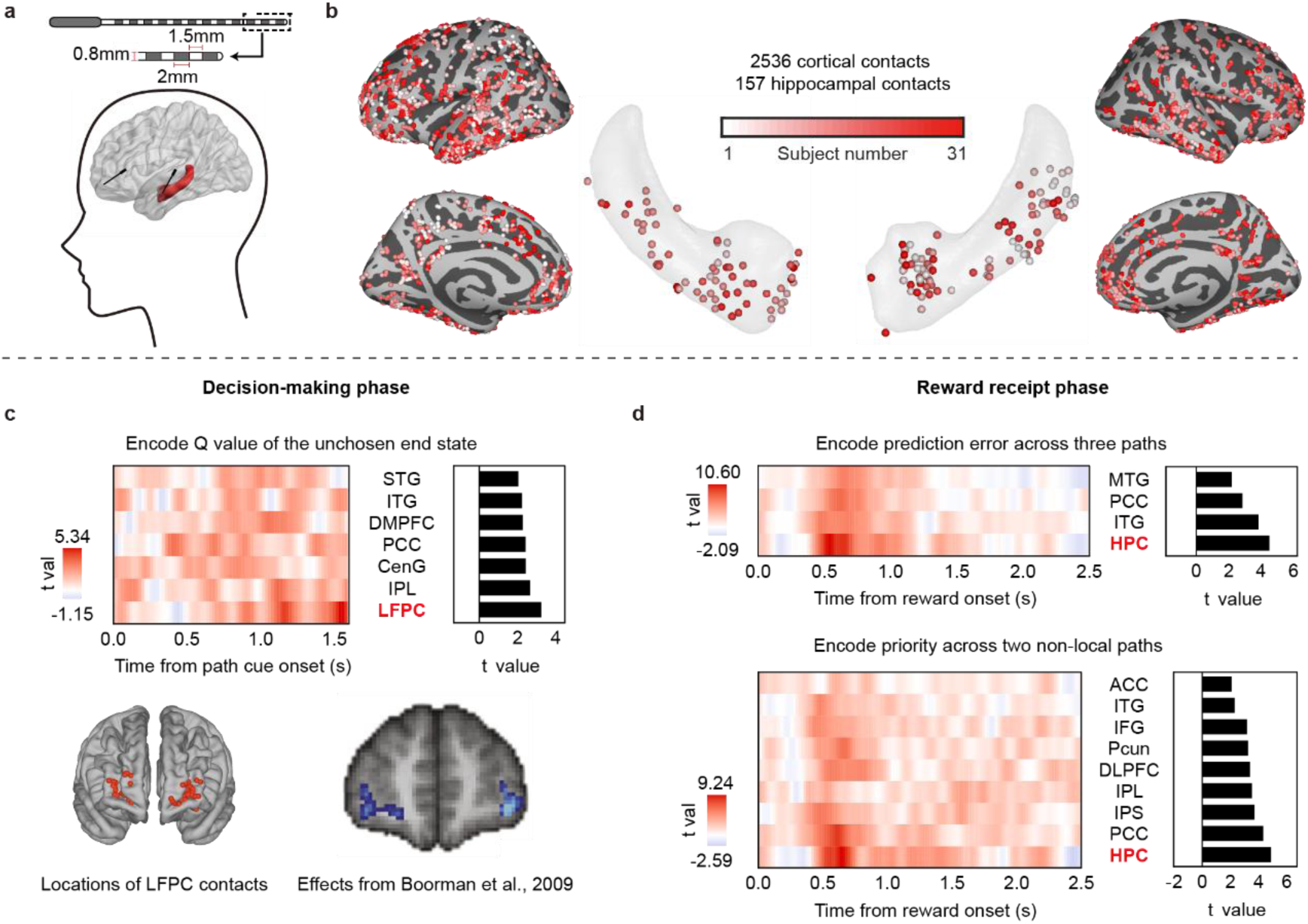
LFPC and hippocampus play crucial roles during decision making and reward processing. **a**, Schematic of intracranial depth electrodes. **b**, Locations of cortical (n = 2536) and hippocampal (n = 157) contacts, colour-coded according to subject ID. **c**, HFB power in the LFPC encoded the value of the unchosen end state during decision-making (from onset of the decision screen - path cue, to the response), consistent with previous fMRI findings (Boorman et al., 2009, 2011). Top: heatmap of t-values (20 ms bins) from a mixed-effects regression of HFB onto unchosen value across ROIs; adjacent bar plot shows the period-averaged t-value. Bottom: anatomical locations of LFPC contacts, and peak voxels reported by Boorman et al. (2009). **d**, During reward receipt (from reward onset to the end of the 2.5 s reward screen), the hippocampus encoded both PE (top) and priority (bottom). Heat-maps (20 ms bins) show t-values from a GLM containing both regressors; bar plots summarise effects over the entire reward period. Colour bars show t-values. Only brain regions showing significant positive effects were shown. ROI abbreviations (used throughout): LFPC (lateral frontopolar cortex), DLPFC (dorsolateral prefrontal cortex), IFG (inferior frontal gyrus), LOFC (lateral orbitofrontal cortex), VMPFC (ventromedial prefrontal cortex), DMPFC (dorsomedial prefrontal cortex), ACC (anterior cingulate cortex), PRE SMA (pre-supplementary motor area), CenG (central gyrus), SPL (superior parietal lobule), IPS (intraparietal sulcus), IPL (inferior parietal lobule), PCC (posterior cingulate cortex), Pcun (precuneus), STG (superior temporal gyrus), MTG (middle temporal gyrus), ITG (inferior temporal gyrus), FuG (fusiform gyrus), VC (visual cortex), HPC (hippocampus).

Although our primary focus is the non-local updating that characterises model-based control, another well-studied hallmark—representation of unchosen option value—can also be tested here (Boorman et al., 2009, 2011). Across the decision period (from path cue onset to choice), we found that the lateral frontopolar cortex (LFPC) robustly encoded the value of the unchosen end state (*β* = 0.014 ± 0.004, *p* = 1.2 × 10^-3^; **Fig. 3c**), replicating prior fMRI findings. Thus, before a choice is made, the LFPC represents alternative values, a signature of model-based evaluation.

To identify regions that update non-local path values at reward, we leveraged our computational model: reward prediction error (PE) should update both local and non-local paths, whereas priority (need × gain) should modulate only non-local updates. For each trial we therefore entered PE (pooled across paths sharing the same end state) and priority (for unseen paths leading to the same end state) simultaneously into a general linear model (GLM) of post-reward HFB. Variance uniquely explained by priority isolates the utility signal guiding prioritised non-local updating; variance explained by PE indexes basic reward surprise. This analysis highlighted the hippocampus, which showed the strongest encoding of both priority (*β* = 0.015 ± 0.003, *p* = 1.1×10^-4^) and PE (*β* = 0.014 ± 0.003, *p* = 6.0 × 10^-4^; both Bonferroni-corrected for all brain areas; **Fig. 3d**) during the 2.5-s reward-receipt period. Together, these results identify the hippocampus and specific prefrontal areas (e.g., LFPC) as key substrates of model-based learning.

### Hippocampal ripples encode the priority of non-local paths

The above results suggested that hippocampal activity at reward carries the integrated priority information needed for model-based learning. We next examined whether hippocampal ripples underpin this learning. Hippocampal ripples, defined as transient high-frequency oscillations, were detected using established methods (A. A. Liu et al., 2022; Norman et al., 2021; Xiao et al., 2025). An example of a ripple waveform and its time–frequency profile is shown in **Fig. 4a**, and the average ripple rate per hippocampal contact appears in **Fig. 4b**.

**Fig. 4.**
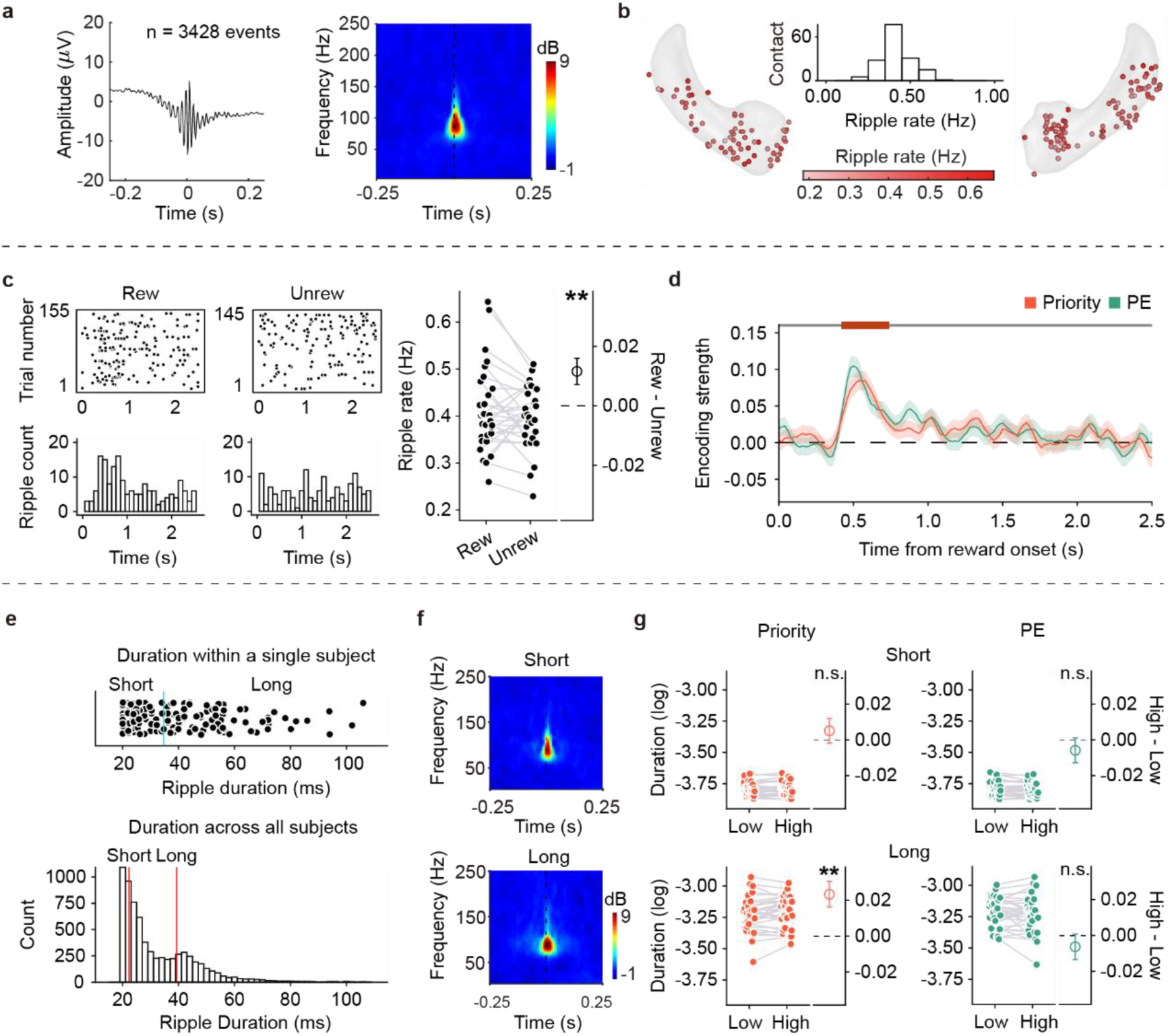
Hippocampal ripples encode priority upon reward receipt. **a**, Example hippocampal ripple event, showing mean field potential (left) and wavelet spectrogram (right) centred on the ripple peak. **b**, Distribution of ripple rates throughout the experiment for all hippocampal contacts included in the analysis. Inset: frequency histogram of ripple rates across contacts. **c**, Ripple rates were higher in rewarded trials than in unrewarded trials during reward receipt. The left and middle panels show data from a representative hippocampal contact with increased ripples following reward delivery. Raster plots (top) and time histograms (bottom) are time-locked to the reward onset. **d**, Hippocampal ripples encoded both priority and PE in the 420–740 ms post-reward interval. Time courses of regression beta weights (20 ms bins) for priority (red) and PE (green) from a single GLM, time-locked to the reward onset. The dark red horizontal bar indicates a significant cluster where hippocampal ripples encoded both priority and PE (cluster-based permutation test). Shaded areas represent SEM. **e**, Top: ripple durations for an example subject. The light blue vertical line indicates the median ripple duration for that subject. Bottom: frequency histogram of ripple durations across subjects. Red vertical lines mark the mean durations of short-(23.062 ± 0.050 ms) and long-duration (42.909 ± 0.248 ms) ripples. Ripples are categorised as short- or long-duration based on the median ripple duration within each subject (short = lower half; long = upper half). **f**, Example wavelet spectrograms of short-duration (top) and long-duration ripples (bottom) from one contact. **g**, Left: In long ripples, ripple duration in trials with high priority was longer than in trials with low priority, whereas this was not the case for short ripples. Trials were median-split by priority in each subject. Right: Ripple durations did not differ between high-PE and low-PE trials in either duration level. Trials were median-split by PE in each subject. In panels **c** and **g**, grey lines connect data from the same subject; each dot is a single subject. Error bars show SEM. n.s.: non-significant; \*\**p* < 0.01.

As both local and non-local learning processes are driven by reward outcome, we first tested whether hippocampal ripples were generally modulated by reward vs. no reward. Indeed, ripple events were more frequent on trials with a reward than trials without reward (*p* = 8.6 × 10^-3^; **Fig. 4c**). This is consistent with animal studies showing ripple occurrence increases following reward delivery (Ambrose et al., 2016; Dupret et al., 2010; Singer & Frank, 2009). Notably, we did not observe a similar reward effect in the hippocampal HFB power (*p* = 0.658). Hippocampal ripples thus appear to be a more sensitive index of reward-related learning than bulk high-frequency power, suggesting that important learning signals may occur specifically during ripple events.

We next tested a specific prediction from our best-fitting RL model: trials starting with rarely seen arms (low-frequency arms) should have a higher priority for non-local updates. Correspondingly, if ripples support prioritised learning, one would expect more ripples in trials starting with a rare arm.

The trial-wise estimation of the model indeed produced systematically different priority values of non-local paths depending on arm frequency (mean priority ± SEM: rare arm trials 0.309 ± 0.010, occasional arm 0.259 ± 0.009, common arm 0.199 ± 0.009). Priority differed significantly across the three arm-frequency levels (*p* = 1.4 × 10^-50^), and was highest for rare-arm trials (rare vs. occasional: *p* = 4.9 × 10^-9^; rare vs. common: *p* < 1 × 10^-50^).

Strikingly, hippocampal ripple rates were also highest in rare-arm trials: there was a significant effect of arm frequency on ripple occurrence during reward receipt (*p* = 6.5 × 10^-10^), driven by more ripples in rare-arm trials compared to occasional (*p* = 2.2 × 10^-5^) or common-arm trials (*p* = 2.3 × 10^-10^). In contrast, during the initial arm presentation (when no reward or value updating occurred), there were no differences in ripple rates between rare vs. other arms (all *p* > 0.05). This rules out the trivial explanation that rare arms simply elicited more ripples due to novelty or greater sensory processing. Instead, the finding suggests that hippocampal ripples are specifically linked to non-local value learning, being most prevalent when the priority for non-local update is highest.

To pinpoint when ripples carry priority-related information, we examined the time course of ripple occurrence after reward. We computed the trial-by-trial ripple rate in 20 ms time bins and regressed it against priority and PE (in the same GLM) as a function of time. This analysis identified a post-reward time window (∼420–740 ms after outcome) during which ripple incidence was significantly associated with both priority and PE (cluster-based permutation test, *p* < 0.01; **Fig. 4d**).

Prior work in rodents indicated that long-duration ripples may carry richer cognitive content (e.g. more comprehensive spatial trajectories) and lead to better memory performance (Fernández-Ruiz et al., 2019). We therefore asked whether long vs. short ripples differed in their sensitivity to priority. Focusing on ripples in the 420–740 ms post-reward receipt window, we split ripple events for each subject at the median duration (on average, short-duration ripple: 23.062 ± 0.050 ms, long-duration ripple: 42.909 ± 0.248 ms; **Fig. 4e, f**).

A mixed-effects ANOVA on ripple duration (log-transformed) (A. A. Liu et al., 2022) revealed a significant modulation by priority for long ripples (high-priority trials elicited longer ripples than low-priority trials, *p* = 1.1 × 10^-3^), whereas short ripples showed no difference (*p* = 0.460; **Fig. 4g**). The interaction between priority level and ripple-length category was marginal significant (*p* = 0.066). In contrast, performing the same analysis with PE did not yield any effect (all *p* > 0.37).

Beyond duration, we also examined ripple amplitude: consistent with the above, priority (but not PE) significantly influenced ripple peak amplitude (higher in high-priority trials) and affected duration distribution overall, whereas neither measure of ripple activity was modulated by PE (see **Extended Data Fig. 3**). These findings indicate that hippocampal ripples – especially longer ripples – are tuned to the priority of forthcoming non-local learning. In other words, beyond simply signalling reward surprise, ripple events also prioritise which non-local experiences to update.

### Cortical reactivation of high-priority path during hippocampal ripple events

Hippocampal ripples are typically accompanied by neural replay (Buzsáki, 2015). A key hypothesis of prioritised model-based learning is that the brain selectively replays high-priority experiences to improve learning efficiency (Y. Liu, Mattar, et al., 2021; Mattar & Daw, 2018). In our task, such a replay event would correspond to neural reactivation of a path image the subject did not actually visit but that shares the same end state following reward receipt. If ripples prioritise non-local learning, reactivation of the high-priority unvisited path should be stronger than that of the low-priority path, and this difference should appear specifically during the ripple.

To test this, we used a neural decoding approach. We first trained one classifier for each of the six path stimuli using stimulus-evoked neural responses (only from cortical contacts) from an independent functional-localiser session, prior to the main task **(Fig. 5a**). We deliberately excluded hippocampal electrodes from this decoding to focus on cortical reinstatement of path representations to avoid any trivial decoding of ripples themselves. We then applied these classifiers to the main task data to detect instances of spontaneous path reactivation.

**Fig. 5.**
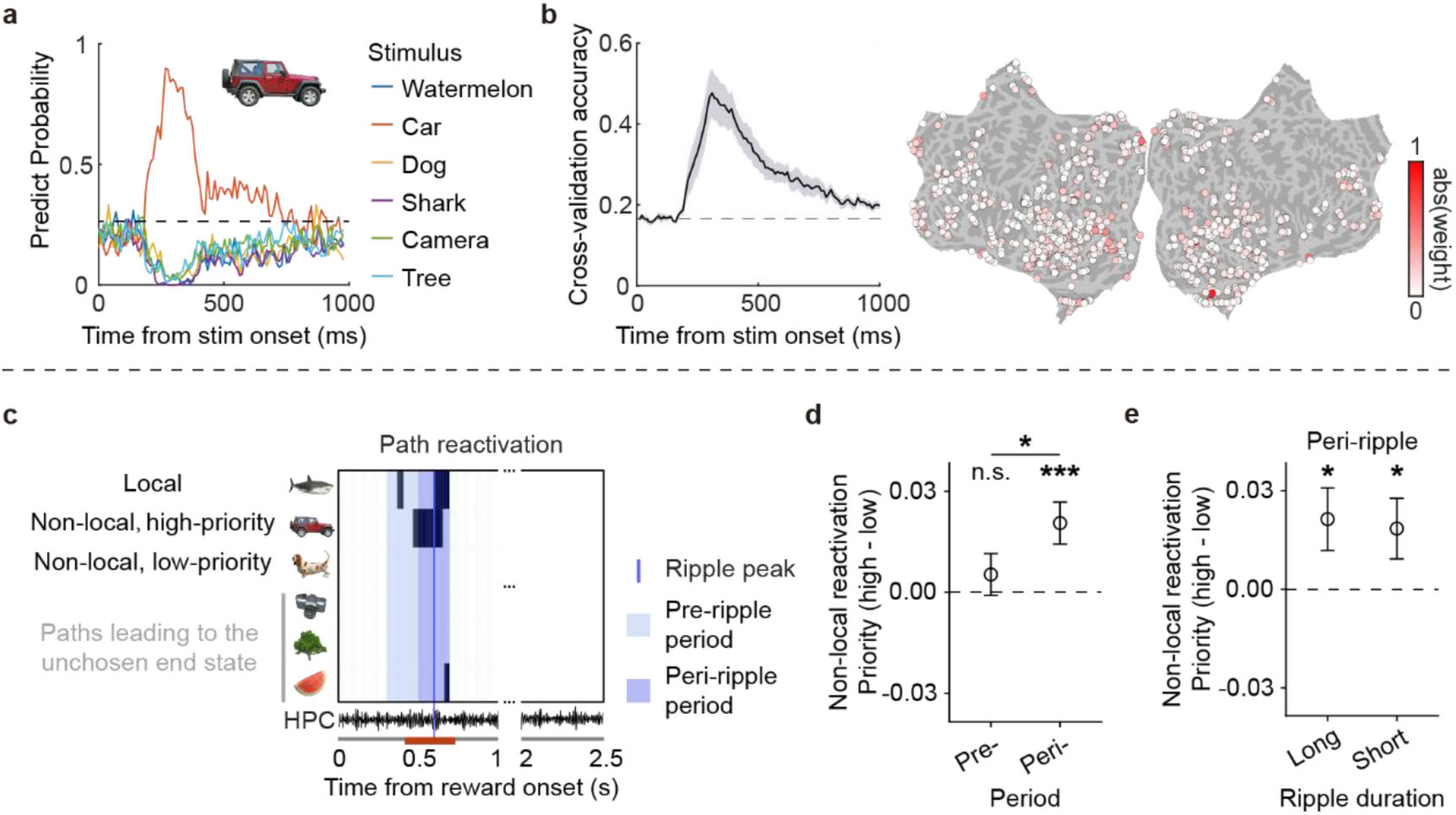
Cortical reactivation of high-priority path during hippocampal ripple period. **a**, Classifier performance from one example subject. Decoding accuracies (in 10 ms bins) for all six classifiers are shown when a specific image was presented in the functional localiser task. The dashed line indicates chance level (label-shuffled data; see Methods). Inset: image presented. **b**, Mean decoding results across subjects. In the functional localiser task, classifiers were trained at each time point and tested across all time points from image onset to 1000 ms post-onset (10 ms bins). Left: decoding accuracy for the 11 subjects whose decoding performance exceeded chance level (label-shuffled; see Methods). Only the accuracies for which classifiers were trained and tested at the same time points were shown. Shaded areas show SEM; the dashed line indicates chance level. Right: absolute classifier weights averaged across all images. Colour bar reflects weight magnitude. **c**, Example trial showing that reactivation of the high-priority non-local path was enhanced in the peri-ripple period compared to the pre-ripple period, whereas reactivation of the low-priority non-local path was not affected. Pre-ripple period: –300 to –100 ms relative to ripple peak; peri-ripple period: –100 to +100 ms. Black regions mark time points where reactivation probability of path stimulus > 0.85; dark-red bar marks ripple cluster encoding both priority and PE. **d**, Reactivation differences between high- and low-priority non-local paths were greater during the peri-ripple period than the pre-ripple period. Non-local path reactivations diverged in the peri-ripple period, whereas no such differentiation occurred in the pre-ripple period. Only ripples whose peaks fell within the 420–740 ms post-reward interval were included in this analysis. **e**, Reactivation differences between non-local paths during the peri-ripple period, shown separately for long-duration and short-duration ripples. The analyses in **d**, **e** controlled for arm frequency and prediction error of non-local path. Error bars show SEM. n.s.: non-significant, * *p* < 0.05, *** *p* < 0.001.

Out of 31 subjects, 24 had completed the localiser session, and 11 of those showed decoding performance above chance (permutation test) for the stimuli (mean cross-validated accuracy 0.477 ± 0.059; **Fig. 5b**; see also **Extended Data Fig. 4a**). We focused on these 11 subjects for the replay analysis. For each subject, we identified the latency at which decoding accuracy peaked during the localiser (on average ∼310 ms after image onset), and then we fixed the decoder at that subject-specific time point for analysing reactivation events in the RL task (see Methods for details).

Using these subject-specific decoders, we quantified the reactivation strength of each path following reward delivery in the RL task. In each trial where the outcome was revealed, there were two non-local paths (the two paths from the other arms that lead to the same end state) that could potentially be updated. We labelled one of these as high-priority and the other as low-priority, based on the trial-by-trial priority values from our winning model (M2). We found that, upon reward receipt, the high-priority unvisited path was reactivated significantly more strongly than the low-priority path (*p* = 0.028). In other words, the neural evidence of replay was biased towards the most important indirect experience. As a control, during the earlier part of the trial (the “arm” screen, before any outcome was revealed), there was no difference in reactivation between the paths that would later be assigned different priority labels. (*p* = 0.107; see also **Extended Data Fig. 4b**). Note, all of the decoding analyses above included arm frequency and PE as covariates, to ensure that differences were specifically related to priority and not to trivial effects of arm novelty or the raw PE signal. This confirms that the priority effect on reactivation emerged specifically during the reward receipt stage, consistent with a prioritised learning mechanism.

Next, we examined the timing of these reactivation events relative to hippocampal ripples. For each ripple that occurred in the critical 420–740 ms post-reward window, we defined a peri-ripple period (–100 to +100 ms around the ripple peak) and a pre-ripple period (–300 to –100 ms before the ripple). We then measured high-priority vs. low-priority path reactivation in these windows (see **Fig. 5c** for a representative example). The high-priority path was reactivated more strongly during the ripple window than before it. In the peri-ripple period, high-priority reactivation was significantly greater than low-priority reactivation (*p* = 4.7 × 10^-4^), whereas in the pre-ripple period there was no difference (*p* = 0.197). Moreover, the magnitude of high–vs– low reactivation difference was itself larger during ripples than before (*p* = 0.040, for peri-vs. pre-ripple comparison; **Fig. 5d**). These results indicate that hippocampal ripples provide time windows that enhance cortical replay of high-priority information. By separating the ripples into long- and short-duration subsets, we observed that the high–low reactivation difference was numerically greater for long ripples, although this trend was not significant (*p* = 0.413; **Fig. 5e**).

These findings indicate that hippocampal ripples gate cortical reactivation of high-priority non-local paths, thereby promoting efficient model-based learning.

### Ripple-aligned LFPC activity underpins non-local learning

Finally, we investigated the role of the prefrontal cortex in orchestrating this prioritised learning, and how it interacts with hippocampal ripples. If a prefrontal region contributes to model-based learning in this task, we reasoned that its involvement should specifically coincide with the hippocampal ripple events that carry priority information. We were especially interested in prefrontal activity time-locked to those ripple events occurring in the 420–740 ms “non-local” window identified above (i.e. ripple events associated with non-local value updates). For comparison, we also examined ripple-aligned prefrontal activity outside that window (defining this as a “control” phase).

We first asked which prefrontal areas showed increased activation during ripple events related to non-local learning (compared to those control ripple events). We found a highly specific engagement of the LFPC: during the non-local ripple phase, LFPC contacts exhibited significantly stronger HFB activity than during the control ripple phase (*p* = 6.2 × 10^-33^; **Fig. 6a**). In other words, whenever a ripple occurred in the critical post-reward window, the LFPC tended to be strongly co-activated, whereas ripple events at other times did not elicit the same LFPC response. This suggests that the LFPC is preferentially recruited in coordination with hippocampal ripples that support non-local learning.

**Fig. 6.**
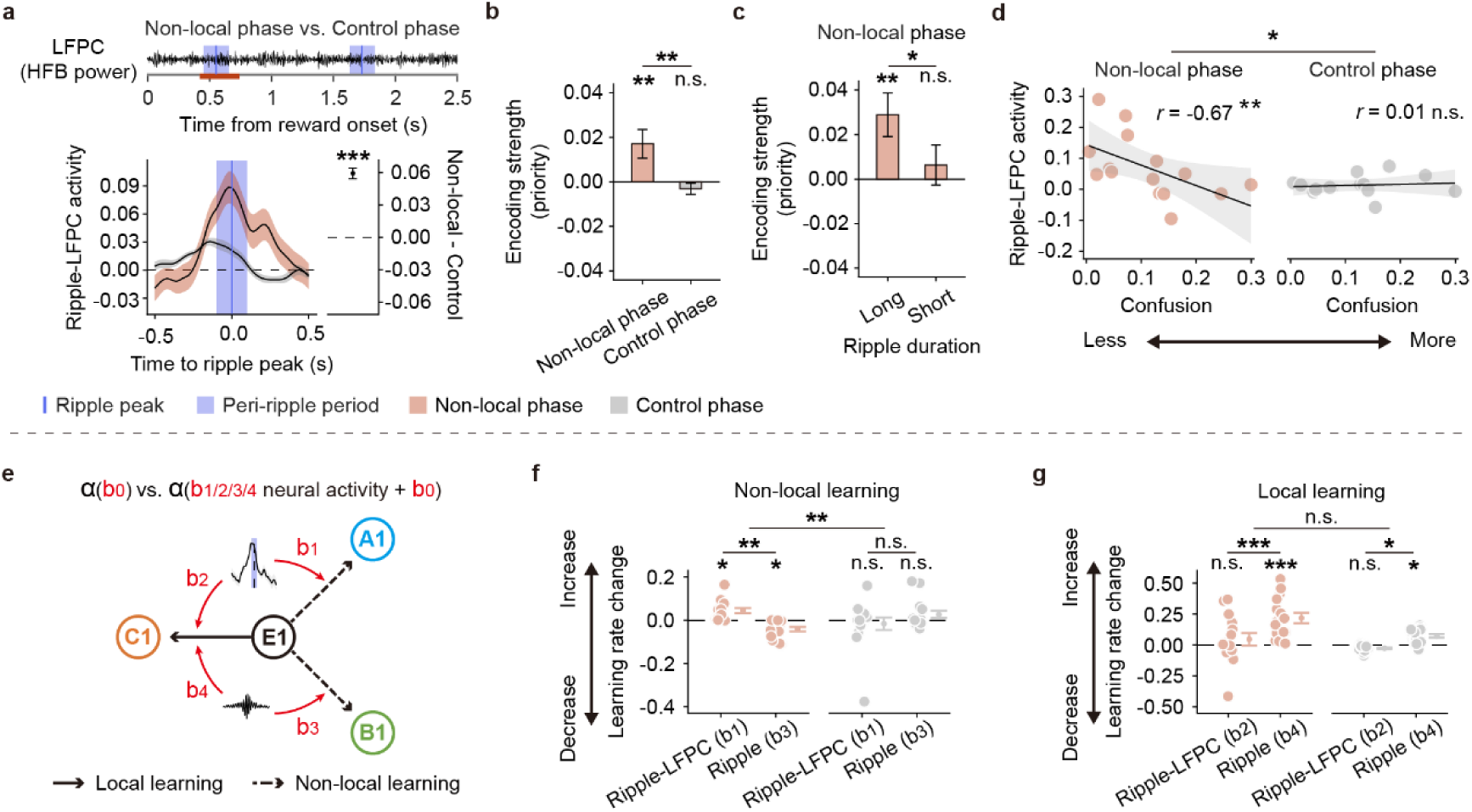
LFPC contributes to non-local learning when aligned to hippocampal ripples. **a**, Peri-ripple LFPC activity was higher during the “non-local” phase than during the “control” phase. The “non-local” phase is defined as peri-ripple events whose ripple peaks fell within the 420–740 ms interval (in dark red) after reward onset, while the “control” phase comprises peri-ripple events with ripple peaks outside this interval. **b**, Peri-ripple LFPC activity encoded priority in the hippocampus-defined “non-local” but not in the “control” phase, and the encoding strength differed between phases. **c**, During “non-local” phase, peri-ripple LFPC activity encoded priority only when aligned with long-duration ripples. This encoding strength was also higher for long-duration ripples than for short-duration ripples. **d**, Peri-ripple LFPC activity during “non-local” phase was negatively correlated with the confusion parameter from our computational model (M2), indicating stronger LFPC activity when value propagation was more accurate (“less confusion”). This relationship was absent during “control” phase, and the correlation coefficient was significantly higher in the “non-local” phase than in the “control” phase. **e**, Illustration of the hybrid model. Each learning rate is the sum of two components. The first component (*b*_1/2/3/4_) captures the contribution of neural activity (peri-ripple LFPC activity or ripple rates alone) to the learning rate; the second component (*b*_0_) serves as an intercept, capturing learning ability not explained by neural activity. If neural activity does not contribute (*b*_1/2/3/4_ = 0), learning rates reduce to the intercept-only model (the same as the “local– non-local” learning model used in Fig. 2c). The *b*_1_, *b*_3_were free parameters for non-local learning, and analogous free parameters (*b*_2_, *b*_4_) were included for local learning. **f**, Peri-ripple LFPC activity explained non-local learning during “non-local” phase but not during “control” phase. **g**, Ripple rates alone (but not peri-ripple LFPC activity) explained local learning during both “non-local” and “control” phases. Error bars represent SEM. Each dot represents a single subject. n.s.: non-significant, * *p* < 0.05, ** *p* < 0.01, *** *p* < 0.001.

We next examined the information content of the LFPC during those coordinated events. Notably, LFPC activity encoded the priority of the upcoming non-local update, but only during the ripple-aligned “non-local” phase. During ripple events in the 420–740 ms window, LFPC signals showed a significant encoding of the priority regressor (*β* = 0.017 ± 0.006, *p* = 7.8 × 10^-3^). However, during ripple events outside this window (“control” phase), the LFPC carried no priority signal (*β* = −0.003 ± 0.002, *p* = 0.216), and the difference in encoding between phases was significant (*p* = 3.4 × 10^-3^; **Fig. 6b**, see also **Extended Data Fig. 5a**). Furthermore, when we looked at LFPC activity outside of any ripple alignment (i.e. general LFPC signals that were not coupled to a hippocampal ripple at all), there was no priority encoding whatsoever (*β* = −0.004 ± 0.007, *p* = 0.614). These results indicate that the LFPC carried prioritised value information exclusively when it was networked with the hippocampus during ripple events.

We also observed an interesting temporal specificity related to ripple duration. If long ripples carry more complex model-based computations (as suggested by our earlier results), one might expect the LFPC to be preferentially engaged by long ripples. Indeed, when ripple events in the “non-local” phase were separated by duration, the LFPC encoded priority significantly during long ripples (*β* = 0.029 ± 0.010, *p* = 3.2 × 10^-3^) but not during short ripples (*β* = 0.006 ± 0.009, *p* = 0.483), and the difference between these was significance (*p* = 0.045, **Fig. 6c**). Thus, long-duration ripples were particularly effective at recruiting the LFPC for prioritised learning.

Throughout all these analyses, we found no evidence that the LFPC encoded PE during the non-local ripple events (*β* = 0.002 ± 0.007, *p* = 0.775). In fact, the LFPC’s priority encoding during ripple events was significantly stronger than any putative PE encoding (*p* = 0.038 for priority vs. PE regression coefficient in the ripple window). By contrast, two other prefrontal regions – the ventromedial PFC (VMPFC) and lateral orbitofrontal cortex (LOFC) – did show the opposite pattern: during ripple events, VMPFC/LOFC activity was modulated by reward PEs (stronger in the non-local phase than control phase, both *p* < 7.4 × 10^-9^) and not by priority (no difference between the non-local phase and control phase, both *p* > 0.05; see also **Extended Data Fig. 5a**), with PE encoding in those regions exceeding any priority signals (both *p* < 9.6 × 10^-6^ for PE vs. priority in non-local phase). This double dissociation underscores that the LFPC plays a unique role in prioritised model-based learning, while other prefrontal areas are more involved in processing the immediate reward outcomes.

If LFPC–hippocampal interactions truly support non-local learning, one would expect that individuals who more effectively engage this circuit should exhibit better model-based performance. Our computational model M2 provided an individual “confusion” parameter χ (higher χ indicates poorer usage of structure). Indeed, we found that subjects with lower confusion (better structure use) tended to have stronger LFPC engagement during ripple events. Specifically, the magnitude of LFPC HFB activity in the non-local ripple phase was negatively correlated with χ across subjects (r = –0.67, *p* = 8.1 × 10^-3^; **Fig. 6d**). This correlation was absent for the control-phase ripple events (r = 0.01, *p* = 0.985; see also **Extended Data Fig. 5b**) and significantly different between the two phases (*z* = –2.04, *p* = 0.041). In other words, when and only when ripple events carried non-local information, greater LFPC activity predicted more accurate generalisation of knowledge. Importantly, this relationship was specific to LFPC: the confusion parameter did not correlate with hippocampal ripple rates or HFB power, nor with VMPFC/LOFC activity, during those same windows (all *p* > 0.25). It was also specific to ripple-aligned LFPC activity. The LFPC signals occurring outside of ripple events showed no such correlation (r = –0.23, *p* = 0.419). These findings indicate that the LFPC’s involvement in non-local learning depends on its synchrony with hippocampal ripples, and that this coordinated activity is directly linked to a participant’s effective use of their internal model.

Lastly, we asked whether the coordinated hippocampal–LFPC activity during ripples had a direct impact on learning itself. If the ripple-coupled LFPC events are truly facilitating non-local learning, then when those events are stronger, the learning rates for non-local updates should effectively be higher. We constructed a hybrid learning model to test this: the model allowed each subject’s (non-local or local) learning rate to be modulated up or down by neural activity on a trial-by-trial basis (see also **Extended Data Fig. 6** for model simulation and parameter recovery). We built two such models – one where the modulator was the LFPC’s peri-ripple HFB activity, and one where it was the hippocampal ripple rate alone – and asked whether including these terms improved the fit over a baseline model with no neural modulators (**Fig. 6e**).

Consistent with our hypothesis, we found that only the LFPC term improved non-local learning during the 420–740 ms ripple phase. There was a significant interaction between neural predictor and phase (*p* = 1.1 × 10^-3^): in the 420–740 ms ripple phase, adding LFPC activity led to a significantly higher non-local learning rate (*p* = 0.021), indicating facilitation of learning, whereas adding hippocampal ripple rate alone actually resulted in a slight decrease in the non-local learning rate (*p* = 0.032; difference between LFPC vs. ripple predictors: *p* = 2.3 × 10^-3^; **Fig. 6f**). In the control phase (ripples outside the 420–740 ms window), neither LFPC nor ripple activity had a significant effect on non-local learning (both *p* > 0.15).

In contrast, for local learning (direct path updates), the hippocampal ripple rate did have a beneficial effect regardless of phase – consistent with the idea that ripple-related replay can also aid memory for directly experienced events – whereas LFPC activity did not influence local learning. Specifically, ripple rate alone significantly increased the local learning rate in both the non-local phase (*p* = 3.76 × 10⁻⁸) and control phase (*p* = 0.039), while adding LFPC activity had no impact (both *p* > 0.18; **Fig. 6g**). These results reinforce a double dissociation: hippocampal ripples on their own support simple, direct learning, but only the combined hippocampus–LFPC circuit supports complex, model-based (non-local) learning. Together, our neural findings indicate that coordinated hippocampal–LFPC activity during ripple events forms a core mechanism for model-based value learning.

## Discussion

Our results reveal a coordinated hippocampal–frontopolar circuit that allows the human brain to learn optimally, extracting the generalisable knowledge from sparse experience. Hippocampal ripples not only signal prediction errors (PEs) but also tag the priority of unvisited options. Working in synchrony with the lateral frontopolar cortex (LFPC), these ripples prioritise access to indirect experiences, driving efficient non-local learning.

Prioritising access to experiences allows the brain to optimise learning by placing high-value information first in line. In our task design, whenever learning – whether local or non-local – was needed, a PE signalled the opportunity for learning to occur (Y. Liu, Mattar, et al., 2021; Mattar & Daw, 2018). We ensured a steady stream of PEs by having reward probabilities drift over time. As expected, hippocampal ripples were sensitive to PE signals at the moment of reward: ripple events were more likely to occur when an outcome was rewarding (positive PE) than when it was not. This finding aligns with rodent studies in which ripples (and associated replay content) are triggered by reward delivery (Ambrose et al., 2016; Foster & Wilson, 2006), and often coincide with phasic activity of reward-responsive neurons in the ventral tegmental area (Gomperts et al., 2015; Kleinman & Foster, 2025).

Beyond responding to reward itself, hippocampal ripples also signalled the priority of nonlocal experience. By analysing priority and PE together, we determined that ripple-related activity encoded the priority of a non-local path over and above what could be explained by the raw prediction error. Notably, we observed that ripple duration was modulated by priority but not by PE: ripples lasted longer on trials where the upcoming non-local value update was high-priority, whereas ripple duration did not systematically vary with the size of the prediction error alone. In our task, priority is a composite quantity – incorporating the reward PE as one component, but also factors like how rarely that path is encountered (need). This observation is consistent with evidence from rodents that long-duration ripples carry more detailed, task-relevant information (Fernández-Ruiz et al., 2019). By extending the ripple, the hippocampus may be able to encode not just that a reward occurred, but also why that reward matters for the broader cognitive map, effectively computing a weighted “priority” for updating other locations.

In line with this functional role of priority, we found that hippocampal ripples biased the reactivation of high-priority information in cortex. During ripple events, the task-related cortical patterns corresponding to the most valuable unvisited path were replayed. This prioritised replay is in agreement with previous human MEG results showing preferential propagation of value along important trajectories (Y. Liu, Mattar, et al., 2021). It also echoes elegant rodent studies: for example, when a rat receives a reward, ripple-associated replay tends to include an alternate route leading to that reward location, if that alternate route has high potential value (Igata et al., 2021; Ólafsdóttir et al., 2015). By providing a temporal “window” after reward during which the most relevant alternative path is reactivated and updated, hippocampal ripples mediate a form of prioritised credit assignment for non-local learning.

Model-based learning further requires that the agent accurately applies its internal model to generalise experiences. In our task, this meant knowing the correct mapping of arms to end states (task structure). We found that the LFPC, a high-level prefrontal region (Amiez et al., 2023; Semendeferi et al., 2001), played a critical role in this process. When non-local learning was occurring in the post-reward period, the LFPC’s involvement correlated each subject’s behavioural use of the task structure: subjects who had learned the structure better (lower confusion) showed stronger LFPC activation during those ripple windows. This relationship was specific in time (it vanished outside the ripple-aligned window) and in region (it was not seen in other frontal areas or in the hippocampus itself). In contrast, we observed no such correlation in the VMPFC or LOFC, two regions often linked to state representation (Constantinescu et al., 2016; Doeller et al., 2010; Schuck et al., 2016) and value processing (Baram et al., 2021; Boorman et al., 2016; O’Doherty et al., 2003; Tom et al., 2007). Consistent with their known roles in value, the VMPFC and LOFC in our study primarily encoded the PE during ripple events, and notably did not carry priority signals in that context. The LFPC, on the other hand, showed the opposite profile: it encoded priority but not PE, and only did so when it was engaged concurrently with a hippocampal ripple. When a ripple was not present, the LFPC showed no priority encoding. These findings underscore that the LFPC, together with hippocampal ripples, is crucial for model-based learning, whereas other frontal regions may contribute more to processing immediate rewards or state values.

The LFPC has been implicated in maintaining internal, goal-relevant information to facilitate exploration and planning (Mansouri et al., 2017). For example, neurophysiological and lesion studies suggest the frontopolar cortex helps evaluate alternative options and switch strategies when needed (Boorman et al., 2009, 2011; Daw et al., 2006; Mansouri et al., 2015). In our study we also saw the LFPC holding the values of unchosen alternatives during choices, replicating previous fMRI results (Boorman et al., 2009, 2011). Our results extend this understanding by showing that LFPC–hippocampal ripple interactions are required for even more complex internal computations, that is prioritising non-local learning. Notably, a fMRI study in primates also found coordinated activation of the hippocampus and frontopolar cortex when monkeys had to use internal model knowledge to guide decisions (Miyamoto et al., 2018). This points to a conserved circuit where the frontopolar cortex interfaces with the hippocampal memory system to support higher-order cognition.

The tight coupling between the hippocampus and LFPC during ripples raises the possibility that LFPC might actively influence the learning process when aligned ripples occur. Indeed, we observed a clear functional impact of LFPC–hippocampal coupling: when the LFPC was strongly active during ripple events, subjects learned more from the reward (higher non-local learning rates), whereas ripple events without LFPC engagement were actually less effective for non-local learning. Consistent with this, monkeys with frontopolar cortex lesions can learn habitual, incremental associations normally but are impaired at one-shot generalisation (Boschin et al., 2015), essentially a deficit in the sort of model-based inference that our task requires. These observations suggest that the precisely time-locked coordination of LFPC activity with hippocampal ripples is the mechanism by which this circuit contributes to model-based learning.

While we uncover a novel hippocampal–LFPC mechanism for model-based learning in humans, there are also limitations to be acknowledged. First, the task required that all three arms converged onto the same two end states (to allow us to test non-local generalisation). This design means that every reward outcome simultaneously created both a local PE (for the currently visited arm) and a non-local PE (for the unvisited arms leading to the same end state). As a result, although we saw hippocampal ripples correlating with PE signals, we could not isolate whether those signals reflected a purely model-based prediction error or just the primary reward itself. Future tasks with separable outcome states could help disentangle direct vs. model-based PEs. Second, we characterised how ripple duration and occurrence related to learning efficiency, but we did not interfere with ripples (Fernández-Ruiz et al., 2019; Igata et al., 2021). One intriguing approach for future work would be to selectively disrupt or enhance LFPC activity time-locked to ripples to see if that perturbs model-based learning (Jacobs et al., 2016; Mohan et al., 2020). Conversely, on the hippocampal side, one could test whether experimentally prolonging ripples (e.g. via stimulation protocols that lengthen ripple duration) would boost the learning of high-priority information.

In summary, our findings show that hippocampal ripples constitute a key cellular mechanism that assigns priority during model-based learning in humans. By synchronising with the frontopolar cortex, these ripples provide a mechanistic account of how the brain learns optimally from sparse experience.

## Methods

### Participants

Thirty-four patients with pharmacologically resistant epilepsy took part in this experiment (20 males, 14 females). All underwent iEEG recording for clinical purposes at Sanbo Brain Hospital of Capital Medical University in Beijing, China, and all were included in the behavioural analysis (see **Extended Data Tables 1, 2** for demographic details and electrode contact distributions). Three subjects lacked hippocampal electrode contacts and were excluded from all neural analyses, leaving 31 subjects with simultaneous recordings in both hippocampal and cortical contacts. All experimental procedures received ethical approval from the Ethics Committee at Sanbo Brain Hospital, Capital Medical University (number: SBNK-YJ-2023-002-01).

### Overview of the experimental design

This experiment, adapted from a previous study (Y. Liu, Mattar, et al., 2021), consisted of three tasks (**Fig. 1**): a functional localiser, a model construction, and a three-armed RL task. The order of these tasks was designed to allow an investigation of neural mechanisms underlying model-based learning. The functional localiser, which preceded any experimental manipulations, was used to train neural decoders for detecting spontaneous reactivations of stimuli in the subsequent RL task. The model construction task ensured that subjects fully understood the task structure of the three-armed RL task and learned arm frequency. Finally, subjects performed the three-armed RL task, during which the key neural mechanisms of model-based learning were investigated. All tasks were implemented using PsychoPy (Peirce et al., 2019).

### Functional localiser

Six distinct visual stimuli, later serving as “paths” in the RL task (e.g., dog – path A1, shark – path A2), were presented in random order during the localiser. Classifiers were trained on the neural activity these stimuli evoked, enabling us to detect spontaneous reactivation of stimuli during the RL task. The mapping between each stimulus and its corresponding path was consistent within a subject but randomised across subjects.

Seven subjects did not undergo this localiser phase.

Each localiser trial began with a fixation cross displayed for 500 ms, followed by a visual stimulus (e.g., a dog) displayed for 1000 ms. Subjects were instructed to consider the semantic meaning of the stimulus (e.g., “this is a dog”). A text label (e.g., “dog” or “car”) then appeared, and subjects were required to respond “yes” within 1000 ms if the text correctly matched the stimulus. Each stimulus was presented 50 times.

### Model construction task

Understanding the task structure in the three-armed RL task was critical for non-local learning. Subjects therefore completed tasks to learn (i) arm–path mappings, (ii) end state–path mappings. In addition, Subjects completed an arm frequency task to learn the frequency (need) with which each arm appeared.

In the arm–path mappings, each of the three arms was deterministically paired with two paths. In this phase each arm occurred equally often. There were three sub-tasks: a learning block (48 trials), a test-with-feedback block (18 trials), and a test-without-feedback block (18 trials). In the learning block, each trial began with a fixation cross (500 ms), followed by an arm stimulus (1500 ms) and then a paired path stimulus (1500 ms). Afterwards, knowledge was tested. In the test-with-feedback block, each trial started with a fixation cross (500 ms), then an arm stimulus (1500 ms). Two paths were shown, and subjects were asked to select the path associated with the arm in a self-paced manner. Feedback indicated whether the selection was correct; if incorrect, both correct paths were displayed (2000 ms), otherwise a correct signal was shown (1000 ms). In the test-without-feedback block, the procedure was identical to the test-with-feedback block except that responses had a 2-second time limit, and no feedback was given.

In the end state–path mapping, subjects learned how each of six paths connected to two end states. Here, each end state was shared by three different paths (e.g., E1 associated with A1, B1, and C1). Learning block (60 trials) and two testing blocks (24 trials each) were conducted similarly to the arm–path mapping.

Subjects repeated both the arm–path mapping and end state–path mapping tasks until at least one of the following criteria was met: (i) test-without-feedback accuracy exceeded 80%, or (ii) subjects could explicitly report all the correct mappings to the experimenter.

In the arm frequency task, subjects learned the probabilities of trials starting at each arm in the RL task (rare 17%, occasional 33%, common 50%). After being informed of these frequencies, subjects completed 30 “perception” trials to experience the distribution. These were followed by 30 “test” trials in which each arm was shown and subjects indicated its frequency (rare, occasional, common).

### Three-armed reinforcement learning task

Following successful completion of the model construction tasks, subjects performed the main three-armed RL task (300 trials; **Fig. 1d**). Each trial began with a fixation cross (500 ms), followed by one arm (2500 ms), which was pseudo-randomly selected according to its assigned frequency. Next, two path stimuli appeared (e.g., A1 and A2, with their left/right positions randomised), and subjects selected one path in a self-paced manner. The chosen end state was then presented, after which subjects pressed the “space” key to proceed to a reward receipt period (2500 ms) displaying the end state’s value (either ¥1 or ¥0). This design avoided immediate visual offset of the end state stimulus in the reward period. Finally, a jittered ITI (500–700 ms) was shown before the next trial. The reward probabilities for each end state followed independent Gaussian random walks (mean 0, standard deviation 0.025, bounded between 0.25 and 0.75; **Fig. 1c**).

### Intracranial EEG recordings

iEEG signals were recorded with a Blackrock clinical monitoring system at a 2000 Hz sampling rate (band-pass 0.3–500 Hz). The depth electrodes had a diameter of 0.8 mm, a contact height of 2 mm, and 3.5 mm spacing between contact centres.

### Electrode anatomical localisation

For each subject, the pre-implantation T1-weighted MRI scans were segmented into grey and white matter using FreeSurfer v7.3.2 (Fischl, 2012). Post-implantation CT scans were co-registered to the pre-implantation MRI using Brainstorm (Tadel et al., 2011). Three independent raters labelled each contact’s location, and any contact more than 3 mm outside the cortical ribbon was excluded. Anatomical labels were assigned according to the Brainnetome atlas (Fan et al., 2016) in individual space. At the group level, cortical templates were resampled and standardised via SUMA (Saad & Reynolds, 2012). Contacts were displayed on a single cortical template (“fsaverage” or “MNI305”) using iELVis (Groppe et al., 2017).

### Preprocessing of iEEG

Preprocessing was performed with MATLAB 2022b (MathWorks Inc.) and EEGLAB v2022.1 (Delorme & Makeig, 2004). First, channels were excluded after visual inspection, if their voltage values, voltage derivative, or root mean square (RMS) at the 99th percentile exceeded 5 standard deviations above the mean of the rest of contacts within the subject. Line noise at 50 Hz and harmonics up to 250 Hz was removed using a zero-lag linear-phase Hamming-windowed FIR band-stop filter (3 Hz wide). Contacts were organised as bipolar pairs: on the same electrode, cortical contacts to an adjacent contact, hippocampal contacts to a nearby white-matter contact. The signal for each contact was referenced to the first contact of that pair.

### Hippocampal ripples detection

Three out of thirty-four subjects were excluded due to a lack of hippocampal contacts. We identified hippocampal ripples using an established pipeline (A. A. Liu et al., 2022; Norman et al., 2021). Contacts within 3 mm of the hippocampal subfields were selected. These contacts were re-referenced to a nearby white-matter contact to reduce common noise. Signals were band-pass filtered between 70 and 180 Hz (zero-lag linear-phase Hamming-windowed FIR filter), and the instantaneous analytic amplitude was computed with a Hilbert transform. To define a robust threshold for ripple detection, we performed several steps. First, the signals were clipped at 4 standard deviations above the mean. The mean and standard deviations were estimated using the least median of squares. Second, the clipped signal was then squared and smoothed using a Kaiser-window FIR low-pass filter with a 40 Hz cutoff. Last, the mean and standard deviation of the smoothed data were used to establish the threshold for ripple detection. Concretely, candidate ripples were events whose amplitude exceeded four standard deviations (SD). The ripple peak was defined as the trough closest to the peak of ripple-band power, and the ripple duration was the period in which the ripple-band power stayed above two SD. Adjacent events within 30 ms were merged, and events shorter than 20 ms or longer than 200 ms were discarded.

Following the consensus (A. A. Liu et al., 2022), we excluded events coinciding with common artefacts (Fiederer et al., 2016), and pathological activities (i.e., interictal epileptic discharge, and pathological high-frequency oscillation) (Bragin et al., 2010; Gelinas et al., 2016). Retained events for each hippocampal contact were visually inspected in time–frequency space, and contacts with abnormal activity were excluded from further analysis.

### High-frequency broadband power

High-frequency broadband (HFB) power was defined as the mean normalised power in the 60–160 Hz range, which correlates with local neuronal spiking (Parvizi & Kastner, 2018). To account for the 1/f power profile, the iEEG signals were divided into 10 Hz sub-bands between 60 and 160 Hz (zero-lag linear-phase Hamming-windowed FIR filters of order 660). The analytic amplitude of each filtered signal, obtained via a Hilbert transform, was normalised and averaged across sub-bands (Fisch et al., 2009).

### Behaviour Analysis

The task design leverages the fact that each end state was shared by three paths, allowing us to separate non-local learning (updating the values of the other two paths converging on the same end state) from local learning (updating only the chosen path). For example, if a reward was obtained in end state E1 through path A1 (“common” arm, blue in **Fig. 1b**), an agent using local learning updates A1, whereas an agent using non-local learning updates B1 and C1. These strategies produce different behavioural patterns depending on arm transition (same vs. different arm). Specifically, if the next trial starts from the same arm, local learning predicts continued preference for A1, whereas a different arm in the next trial prompts non-local learning to prefer B1 or C1, but not local learning.

Hence, if agents rely on both local and non-local learning, being rewarded in one trial should increase the probability of selecting a path leading to the same end state in the next trial, whether or not the arm changes. Crucially, if the agent fully utilises the learned task structure, the “stay” probability should remain similar regardless of whether the arm is repeated or changed.

We used a mixed-effects logistic regression to measure local and non-local learning in a model-agnostic way. The dependent variable was the choice in each trial, coded as “stay” (choosing the same end state) vs. “switch” (selecting a different end state). The predictors included the last-trial outcome (rewarded vs. unrewarded), whether the arm changed or not on the current trial, and their interaction. The intercepts, main effects, and interactions were specified as random effects (allowed to vary across subjects):

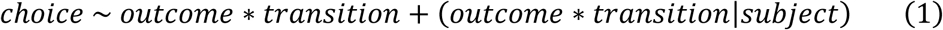

Local learning was defined as the simple effect of the last-trial outcome in the same arm condition, while non-local learning was defined as the simple effect of the last-trial outcome in the different arm condition. The efficiency gap between these two forms of learning corresponded to the interaction between outcome and arm transition.

### Behaviour Modelling

To investigate the computational mechanisms underlying value learning in this task, we built a series of RL models, all based on a baseline model (Y. Liu, Mattar, et al., 2021), that accommodates both local and non-local learning but allows different learning rates for these paths. This baseline model was referred to as the “local-non-local” learning model. In general, the models assume that subjects’ choices are guided by a value function *Q*(*s*, *a*). The choice in each trial is generated by a softmax function:

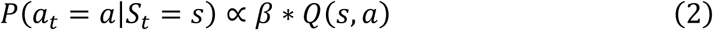

where *β* is the inverse temperature. Here, the action *a* is defined in terms of the end state (E1 or E2) it leads to, and the state *s* refers to the arm (common, occasional, or rare). Consequently, *Q*(*s*, *a*) corresponds to the value of a path associated with (*s*, *a*). In the baseline model, once a reward (*r*) is received by selecting an action, the corresponding *Q* values in all three arms are updated simultaneously in each trial, leading to three learning rules:

Local learning:

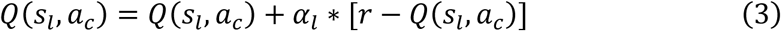

Non-local learning:

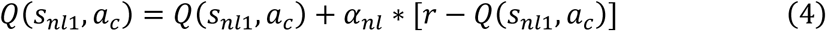

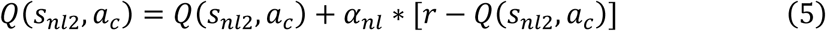

where *s*_*l*_ represents the local arm and *s*_*nl*1_, *s*_*nl*2_ are the two non-local arms, while *α*_*l*_ and *α*_*nl*_ denote the local and non-local learning rates, respectively. Setting *α*_*nl*_ = 0 would reduce the model to a simpler, local-learning-only form. *α*_*c*_ represents the chosen end state.

To test whether non-local value learning was prioritised, we augmented the model by introducing separate learning rates for high-vs. low-level non-local arms. In each trial, the non-local arms are labelled *s*_*high*_ or *s*_*low*_ based on priority. The update rules then become:

Local learning:

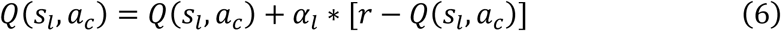

Non-local learning (high level):

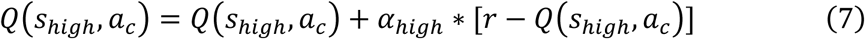

Non-local learning (low level):

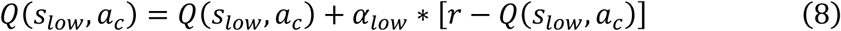

Priority was defined as the product of need and gain (Agrawal et al., 2022). Need refers to the predetermined frequency of each arm, while gain reflects (i) the bellman residual (absolute prediction error), and (ii) the value of visiting a rarely experienced arm (akin to uncertainty). Formally:

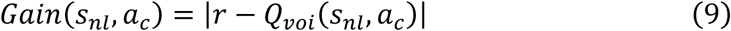

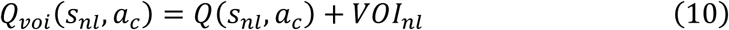

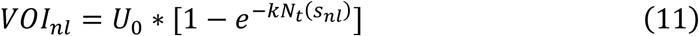

where *s*_*nl*_ indicates a non-local arm, *U*_0_ is the upper bound of the uncertainty, *k* is the hazard rate of mode switching, and *N*_*t*_(*s*_*nl*_) is the number of trials since the last visit to this arm. *N*_*t*_(*s*_*nl*_) is initialised at 10,000, making the *VOI*_*nl*_ close to *U*_0_ at the outset and then dropping to 0 once *s*_*nl*_ is visited. This model was referred to as M1 in the main text.

Finally, to explicitly account for the efficiency gap between local and non-local learning, we introduced “confusion” mechanism, updating alternative paths in unvisited arms:

Confused learning (high level):

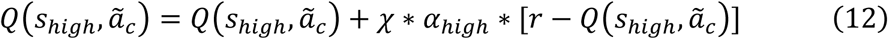

Confused learning (low level):

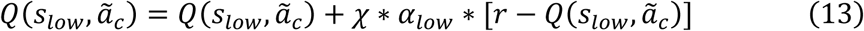

where *χ* measures deficits in ability of structure utilisation, and *ã*_*c*_ leads to the opposite end state. This model was referred to as M2 in the main text.

We were interested in two candidate models: M1 (priority), M2 (priority + confusion). We estimated them hierarchically in Stan (Carpenter et al., 2017), with subject-level parameters drawn from group-level normal priors (*μ* = 0 and *σ* = 1). Each model ran four chains of 2000 samples, and discarded the first 1000 warm-up samples. Visual inspection (**Extended Data Fig. 1**) and R^ < 1.1 (Gelman & Rubin, 1992) confirmed chain convergence. Across both fits, the M2 model better explained participants’ behaviours (**Fig. 2d**).

### Stimuli decoding and reactivation detection

Reactivation of path stimulus was measured to test whether the non-local paths were prioritised. Following our previous work (Y. Liu, Dolan, et al., 2021; Y. Liu et al., 2019; Y. Liu, Mattar, et al., 2021), we trained separate lasso-regularised logistic regressions for each path stimulus (e.g., A1, A2) on pre-processed signals from the functional localiser task. Because we were primarily interested in ripple-aligned neural reactivations, we excluded hippocampal contacts during classifier training to avoid bias from hippocampal signals. Each classifier, using a 10-fold cross-validation scheme, distinguished one stimulus (e.g., A1) from all others. Prediction accuracy was measured by the proportion of test trials where the classifier predicts the correct trial label with the highest probability. Because 7 subjects did not participate the functional localiser tasks, there were 24 out of 31 subjects in this analysis.

Owing to variability in electrode placement across subjects, high classification accuracy could not be guaranteed for each subject, in contrast to previous MEG studies (Y. Liu et al., 2019; Y. Liu, Mattar, et al., 2021). To select subjects with high classification accuracy, we generated a null distribution of classification accuracies by randomly shuffling stimulus labels and retraining classifiers 100 times for each subject. A subject was included in the reactivation analysis if, for every path classifier, the real-label accuracy exceeded the 95th percentile of this null distribution. Following these criteria, 11 subjects were selected (**Fig. 5b** and **Extended Data Fig. 4a**).

Each subject’s classifiers were then trained at the time point yielding highest accuracy. The classifiers were then applied to the pre-processed signals of the three-armed RL task at the corresponding time window, covering the arm screen, reward receipt, and pre-, peri-ripple periods, producing a [time × path] reactivation probability matrix per trial. Pre-ripple (−300 to −100 ms), and peri-ripple (−100 to +100 ms) time windows were defined relative to the ripple peak.

### Mixed-effects analysis

We conducted mixed-effects analyses in R (R Core Team, 2013) using the LME4 package (Bates et al., 2015).

To test reward effect in behaviours, we used a mixed-effects logistic regression. The intercepts, main effects were specified as random effects:

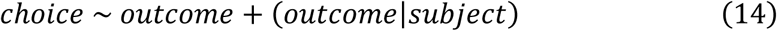

where *choice* was coded as “stay” (choosing the same end state) vs. “switch” (selecting a different end state). *outcome* was a factor variable indicating whether the previous trial was rewarded or not.

To test whether priority differed as a function of arm frequency, we used a mixed-effects logistic regression. The intercepts were specified as random effects:

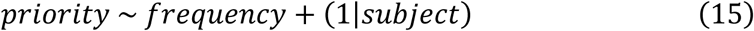

where *priority* was the trial-wise priority for each non-local path estimated by our computational model. *frequency* was a factor variable (rare, occasional, common).

To test whether neural activity contributed to learning ability, we used the following model with subject included as a random intercept:

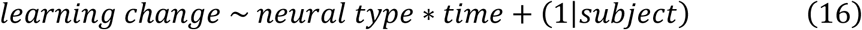

where *learning change* was the difference between *ϕ* (intercept + slope) and *ϕ* (intercept). Both intercept and slope were estimated from the hybrid model. *neural type* and *time* were factor variables. *neural type* included “ripple rate” and “peri-ripple LFPC activity”. *time* included the “non-local” and “control” phases. For all neural analyses, unless otherwise stated, only the intercept was considered a random effect in each model. When dealing with data at the contact level, “contact” was nested within “subject” to reflect our data’s hierarchical structure, that is, (1|subject/contact).

To determine which regions of interest (ROIs) encoded the unchosen-end state value during decision-making, we used the following model:

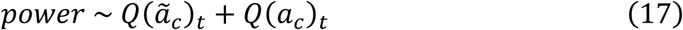

where *power* was the power in high frequency broadband. *Q*(*ã*_*c*_)_*t*_ was the average of the *Q*-values of the three paths leading to the unchosen end state in the *t*-th trial, and *Q*(*α*_*c*_)_*t*_ was the average of the *Q*-values of the three paths leading to the chosen end state. The *Q*-values were averaged due to the high inter-correlation among those paths.

To test whether ROIs encoded priority and prediction error (PE) during reward receipt, we used:

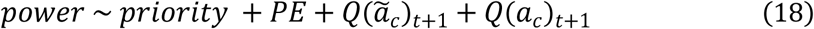

where *priority* was the summed priority of the two non-local paths, and *PE* was the averaged value of *PE* across the three paths converging on the same end state. Due to high collinearity, the priorities of non-local paths were summed and the PEs of the three paths were averaged. The terms *Q*(*ã*_*c*_)_*t*+1_ and *Q*(*α*_*c*_)_*t*+1_ were updated *Q* −values after receiving reward in the *t*-th trial, serving as control variables.

To investigate whether hippocampal ripples encoded reward-related information at reward receipt, we used:

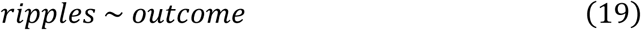

As a control analysis, we reused the structure from Eq. (19), replacing the dependent variable with *power* in the hippocampus.

To examine if hippocampal ripples were involved in prioritised non-local learning during reward receipt, we adopted:

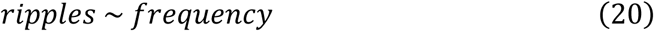

where *frequency* was a factor variable. *frequency* was the arm frequency (rare, occasional, common). Note that our task design produced different priority values depending on arm frequency, but this did not mean that priority was equivalent to frequency. As a control analysis, we reused the structure from Eq. (20), replacing the dependent variable with *ripples* during the arm screen.

To assess whether hippocampal ripples encoded both priority and PE during reward receipt, we conducted two analyses. First, we reused the structure from Eq. (18), replacing the dependent variable with *ripples*.

Second, we tested whether ripple attributes were modulated by priority:

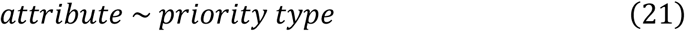

where *attribute* refers to duration, amplitude, and frequency, respectively. *priority type* was high vs. low priority based on median value within subjects. As a control analysis, we reused the structure from Eq. (21), replacing the *priority type* with *PE type*.

We further tested whether long-duration ripples were specifically modulated by priority:

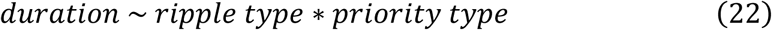

where *ripple type* was long vs. short duration. As a control analysis, we reused the structure from Eq. (22), replacing the *priority type* with *PE type*.

To validate our model’s assumption that non-local learning depended on non-local path priority, we tested whether reactivation strength was predicted by priority type:

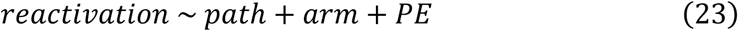

where *path* (high-priority vs. low-priority non-local path), *arm* (common, occasional, or rare) were factor variables, and where *PE* was continuous variable. *arm* and *PE* served as control variables.

To determine whether the difference in reactivation between non-local paths changed by stage during reward receipt, we used:

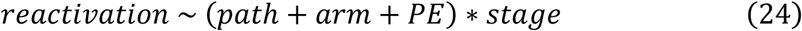

where *stage* (pre-ripple vs. peri-ripple) was factor variables.

We tested whether the difference in reactivation between non-local paths changed by ripple duration during peri-ripple stage:

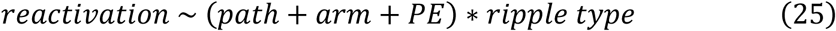

Finally, to test whether peri-ripple HFB power in specific ROI was higher in the “non-local” phase (peri-ripple events whose ripple peaks fell within 420–740 ms) than in the “control” phase (all other peri-ripple events), we used:

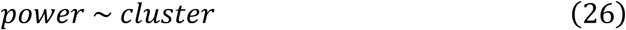

where *cluster* was a factor variable distinguishing the “non-local” phase from the “control” phase.

To examine whether peri-ripple HFB power encoded priority and PE exclusively in the “non-local” phase, we used:

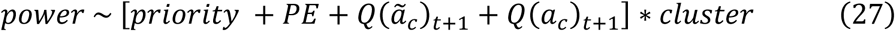

To test whether the priority-related effects were stronger in long-duration ripples, we reused structure from Eq. (27), replacing the *cluster* with *ripple type* and the dependent variable with *power* during “non-local” phase.

### Statistical analysis

Descriptive statistics are presented as mean ± standard error of the mean (Mean ± SEM). Unless otherwise stated, linear mixed-effects models were used for both behavioural and neural data (described in detail above). Significance tests were primarily two-tailed with a threshold of *p* < 0.05. Analyses guided by a strong hypothesis were one-tailed with a threshold of *p* < 0.05, specifically, all reactivation analyses and comparisons of priority-encoding strength in the LFPC between long- and short-duration ripples.

To control for multiple comparisons in the temporal analysis of hippocampal ripples, we used a cluster-based permutation test (Maris & Oostenveld, 2007). In this test, we generated a null distribution by shuffling either time points or trial identity 100 times. Both approaches yielded significant results (*p* < 0.01), and we report the results based on time-point shuffling.

Correlations between confusion and efficiency gap were assessed with Pearson correlation, whereas correlations between confusion and peri-ripple activity used Spearman’s rank correlation. Comparisons between correlation coefficients were performed using Steiger’s approach. Multiple comparisons, excluding those in the temporal analyses, were controlled using Bonferroni correction.

## Acknowledgments

We thank T. Behrens, N. Daw, for helpful discussions; J. Ou, and J. Wang for comments on this manuscript. **Author contributions**: Conceptualization, Y.L., and X.Z.; Investigation, X.Z., X.W., X.H., H.W., Q.Y., L.H., J.Z., Z.X., J.X., and Y.L.; Writing - Original Draft, X.Z., Y.L.; Writing - Review & Editing, X.Z., and Y.L. **Funding**: this study is supported by the National Science and Technology Innovation 2030 Major Programme (2022ZD0205500), the National Natural Science Foundation of China (32271093), the Beijing Natural Science Foundation (Z230010, L222033), and the Fundamental Research Funds for the Central Universities.

## Conflict of interest

The authors have indicated they have no potential conflicts of interest to disclose.

## Data Availability

The processed neural data (i.e., detected ripple events, high-frequency broadband activity and cortical reactivation), and corresponding behavioural and modelling data, will be made available at https://zenodo.org/uploads/15580631 upon publication.

## Code Availability

The analysis code will be accessible at https://gitlab.com/liu_lab/three_arm_seeg upon publication.

## Extended Data Information

**Extended Data Fig. 1.**
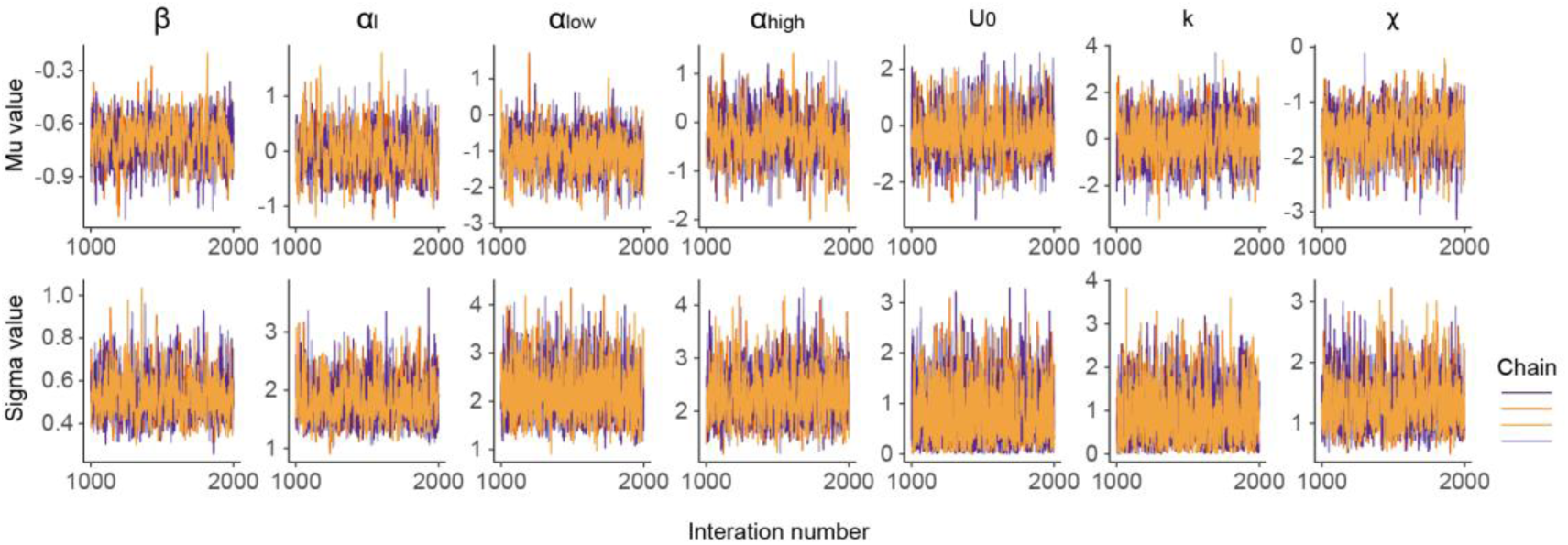
Model convergence. Model fitting is done performed using an MCMC algorithm (Betancourt, 2017). The convergence of four Markov chains for the group-level parameters in the winning model (M2) confirmed reliable estimation. *β* is the inverse temperature. *α*_*l*_ is the learning rate for the local path. *α*_*low*_ is the learning rate for the low-priority non-local path. *α*_*high*_ is the learning rate for the high-priority non-local path. *U*_0_ is the upper bound of the uncertainty. *k* is the hazard rate of mode switching. *χ* is the confusion parameter measuring ability of structure utilisation. The panels in the first row show the means of the parameter distributions in Markov chains, and those in the second row show the standard deviations.

**Extended Data Fig. 2.**
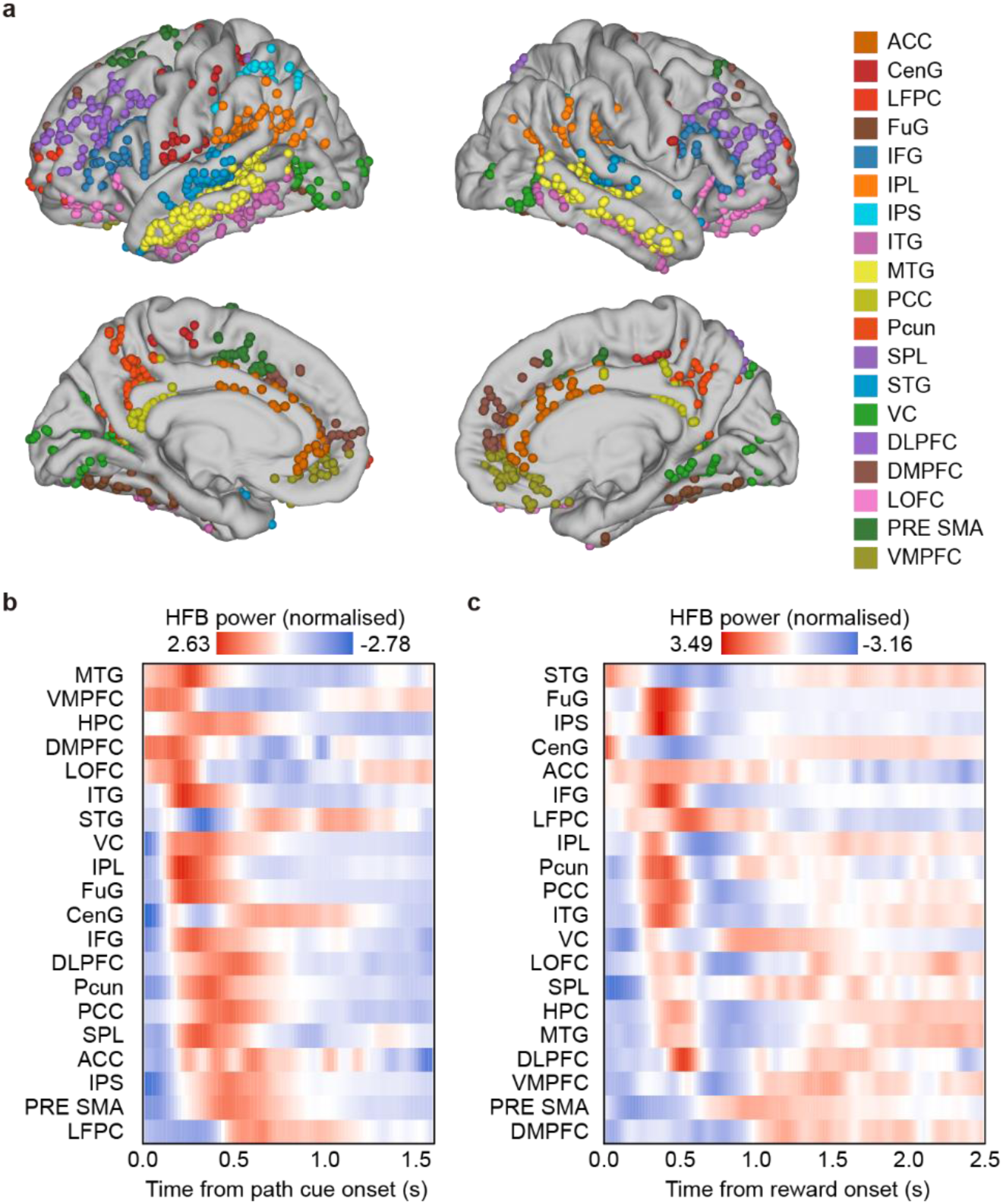
HFB power changes of different brain regions during decision-making and reward receipt. **(a)** Anatomical distribution of cortical contacts coloured based on their respective ROIs. **(b)** The time course of HFB power during decision-making for each ROI, time-locked to the onset of decision-making screen. **(c)** The time course of HFB power during reward receipt for each ROI, time-locked to the reward receipt onset. These regions were sorted by the time at which their neural activity first exceeded zero.

**Extended Data Fig. 3.**
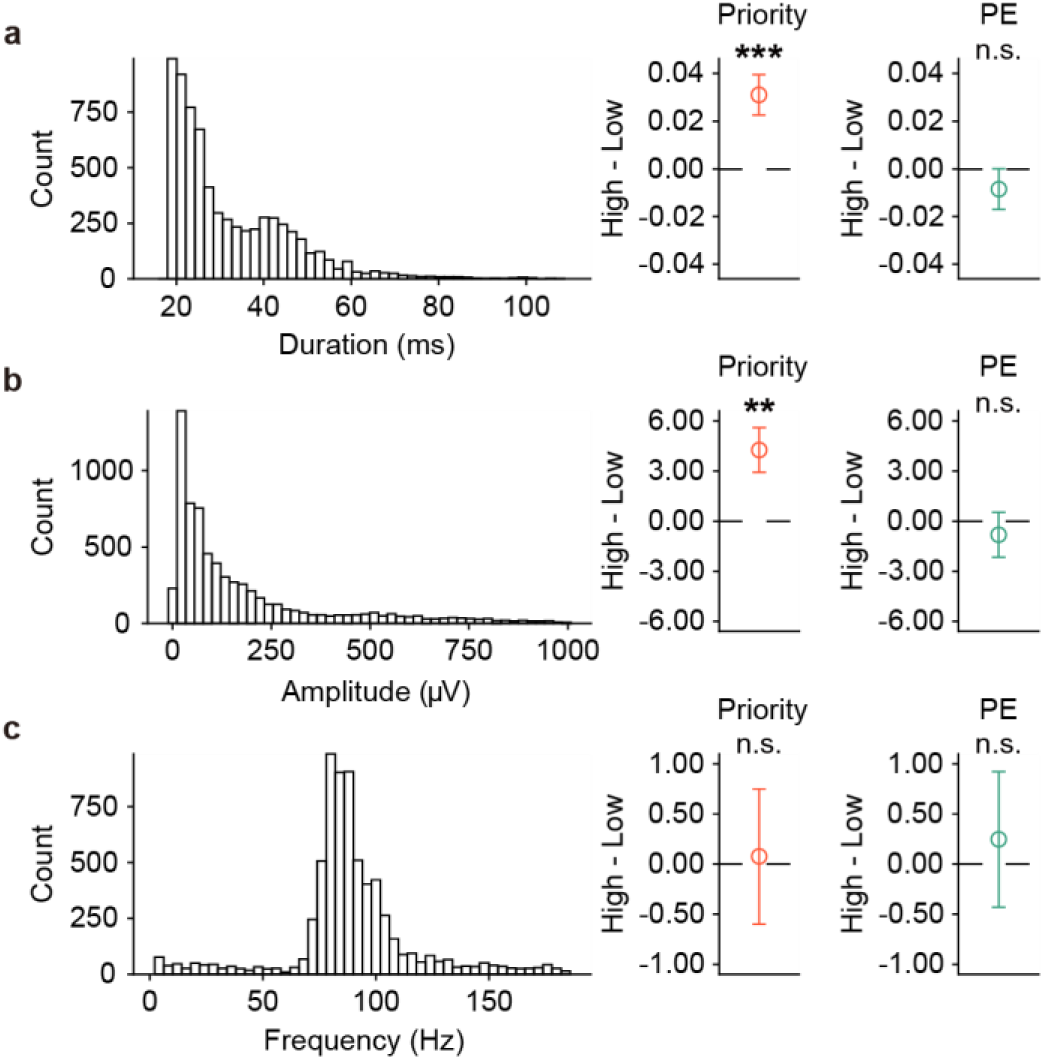
Priority selectively modulated attributes of hippocampal ripples. **(a)** Duration of hippocampal ripples in trials with high priority was longer than in those with low priority, whereas duration was not modulated by prediction error. **(b)** Peak amplitude of hippocampal ripples in trials with high priority was greater than in those with low priority, whereas it was not modulated by prediction error. **(c)** Peak frequency of hippocampal ripples was not modulated by either priority or prediction error. In **a**-**c**, trials were median split by condition (either priority or prediction error) within each subject.

**Extended Data Fig. 4.**
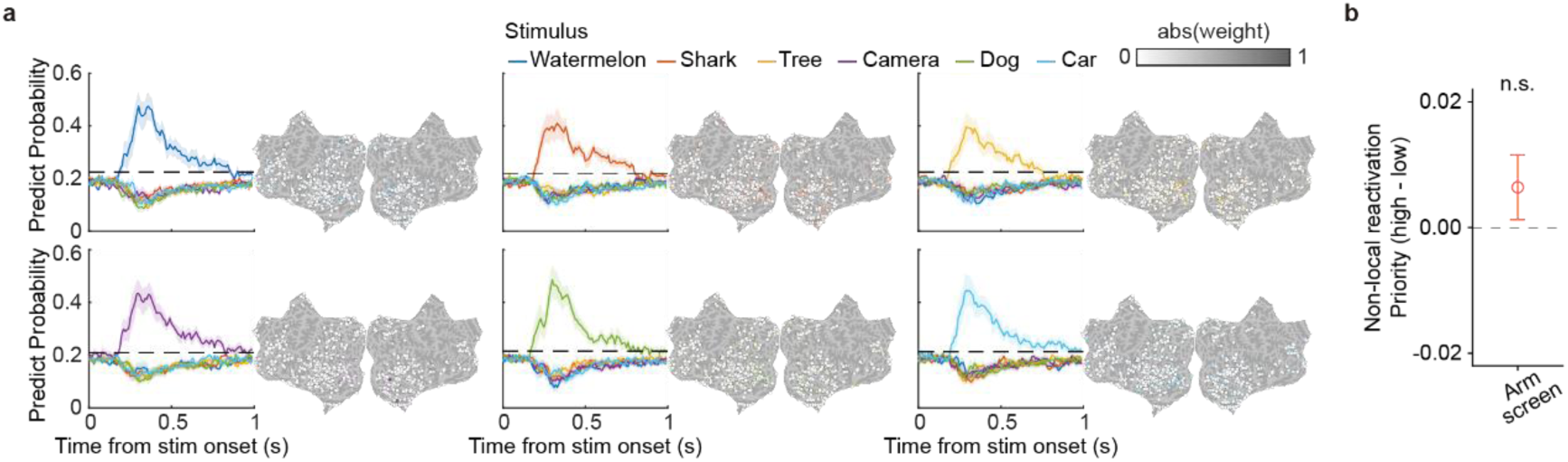
Decoding performances for all stimuli and prioritised reactivation. **(a)** Decoding performance for all six images. In each panel, the left plot shows predicted probabilities from the classifier for a specific image; the right plot shows absolute classifier weights. Performances are averaged across 11 subjects. In the functional localiser session, classifiers were trained at each time point and tested across all time points from image onset to 1000 ms post-onset (10 ms bins) using a ten-fold cross-validation scheme. Only probabilities for classifiers trained and tested at the same time points are shown. The dashed line denotes chance level (label-shuffled data). Shaded areas indicate SEM. Colour bar reflects weight magnitude. **(b)** Reactivation strength showed no difference between the high- and low-priority non-local paths during the arm screen. This analysis controlled for arm frequency and prediction error. Error bars show SEM. n.s.: non-significant.

**Extended Data Fig. 5.**
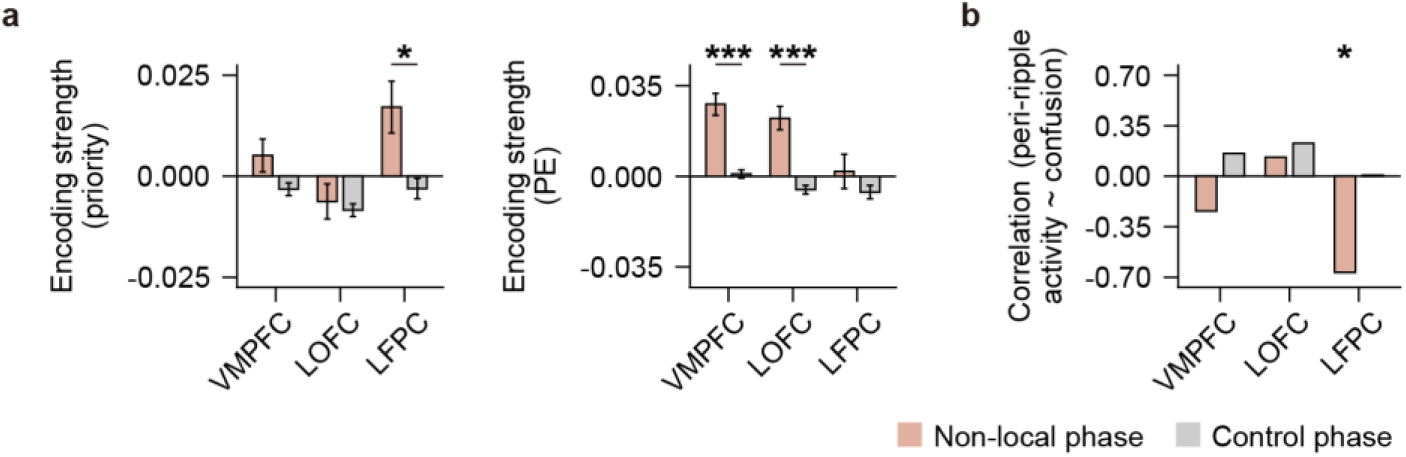
Contribution of different prefrontal areas to learning when aligned to hippocampal ripples. **(a)** In the LFPC, peri-ripple HFB activity encoded priority more strongly in the “non-local” phase than in the “control” phase. In the VMPFC and LOFC, peri-ripple HFB activity encoded PE more strongly in the “non-local” phase compared to the “control” phase. **(b)** Among all prefrontal areas tested, only the peri-ripple LFPC activity correlated with the confusion parameter from the winning computational model, and this correlation was specific to the “non-local” phase. Statistical values reported in this figure were Bonferroni-corrected for the three areas. * *p* < 0.05, *** *p* < 0.001.

**Extended Data Fig. 6.**
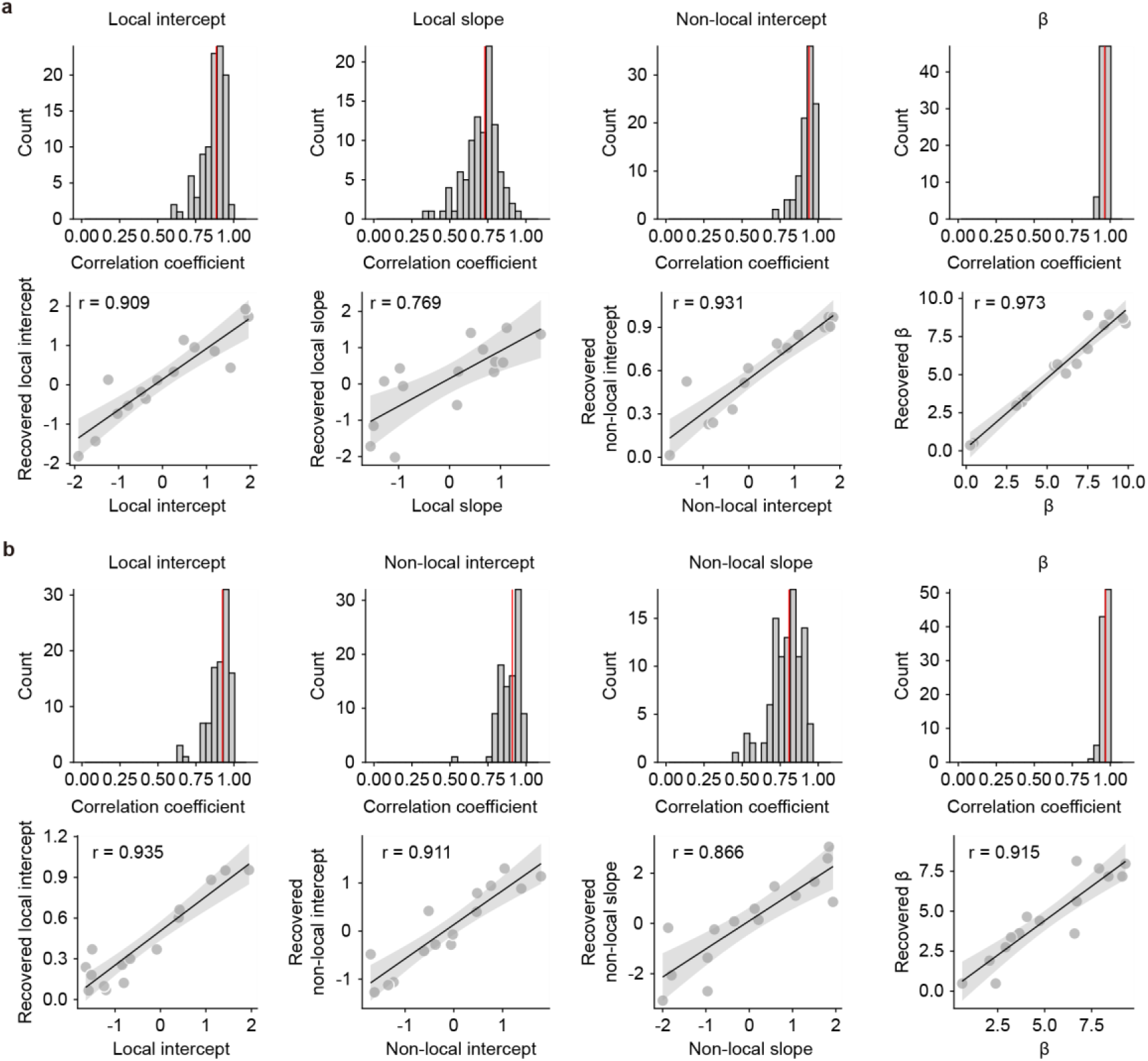
Validation of the hybrid models. **(a)** Free parameters in the local hybrid model were well recovered. The top panel shows the distribution of parameter recovery. The x-axis indicates the correlation between recovered values and true values. The red vertical line indicates the mean value of the distribution. And the bottom panel presents an example recovery. In each recovery, free parameters for 15 subjects were randomly generated. Behaviours were then simulated according to those parameters and the hybrid model, after which recovered parameters were estimated by fitting the model to the simulated behaviours. This procedure was repeated 100 times. In this model, the local learning rate is defined as the sum of the local intercept and local slope after *ϕ* transformation, whereas the non-local learning rate is defined as the non-local intercept after *ϕ* transformation. Here, *ϕ*(*x*) denotes the standard normal cumulative distribution function. **(b)** Free parameters in the non-local hybrid model were also well recovered. The top panel again shows the distribution of parameter recovery, and the bottom panel provides an example recovery. The recovery procedure was identical to that used for the local hybrid model. In this model, the non-local learning rate is defined as the sum of the non-local intercept and non-local slope, while the local learning rate is defined as the local intercept. In both models, *β* is the inverse temperature.

**Extended Data Table 1.** Demographic information of patients.

| Subject # | Gender | Handedness | Age | Education years | RPM score | MMSE |
| --- | --- | --- | --- | --- | --- | --- |
| SUB01 | male | right | 28 | 16 | 17 | 26 |
| SUB02 | male | right | 11 | 9 | 41 | 25 |
| SUB03 | male | right | 15 | 9 | 58 | 30 |
| SUB04 | male | right | 36 | 12 | 32 | 27 |
| SUB05 | male | right | 12 | 9 | 15 | 30 |
| SUB06 | female | right | 30 | 16 | 49 | 27 |
| SUB07 | male | right | 22 | 14 | 41 | 30 |
| SUB08 | male | right | 15 | 9 | 23 | 28 |
| SUB09 | female | right | 37 | 16 | 36 | 30 |
| SUB10 | female | right | 23 | 16 | 55 | 30 |
| SUB11 | male | right | 19 | 13 | 47 | - |
| SUB12 | male | left | 24 | 16 | 47 | 30 |
| SUB13 | male | right | 18 | 12 | - | 29 |
| SUB14 | male | right | 19 | 12 | 56 | 30 |
| SUB15 | male | right | 30 | 12 | 52 | 30 |
| SUB16 | female | right | 14 | 8 | 52 | 30 |
| SUB17 | male | right | 24 | 15 | 49 | 26 |
| SUB18 | female | left | 26 | 19 | 50 | 30 |
| SUB19 | male | right | 15 | 9 | 54 | 30 |
| SUB20 | male | right | 16 | 10 | - | 30 |
| SUB21 | male | right | 26 | 16 | 55 | 30 |
| SUB22 | male | right | 27 | 16 | 56 | 30 |
| SUB23 | female | right | 54 | 16 | 44 | 30 |
| SUB24 | male | right | 37 | 16 | 31 | 30 |
| SUB25 | female | right | 33 | 9 | 49 | 28 |
| SUB26 | male | right | 30 | 16 | 42 | 29 |
| SUB27 | female | right | 30 | 12 | - | 29 |
| SUB28 | female | right | 51 | 16 | - | 30 |
| SUB29 | female | right | 32 | 16 | - | 30 |
| SUB30 | male | right | 26 | 9 | - | 27 |
| SUB31 | female | right | 29 | 16 | - | 30 |
| SUB32 | female | right | 24 | 12 | - | 27 |
| SUB33 | female | right | 25 | 9 | - | 28 |
| SUB34 | female | right | 37 | 9 | - | 30 |
| M±SE | 14/34 | 2/34 | 26.3±1.7 | 12.9±0.6 | 43.8±2.5 | 29.0±0.3 |

Note. The RPM score is the raw score of Raven's Progressive Matrices, and the MMSE is the raw score of the Mini-Mental State Examination. '-' indicates missing data for the item.

**Extended Data Table 2.** The number of contacts in each ROI for each subject.

| Subject # | HPC | LFPC | CenG | FuG | IFG | IPL | IPS | ITG | MTG | PCC | Pcun |
| --- | --- | --- | --- | --- | --- | --- | --- | --- | --- | --- | --- |
| SUB01 | 8 | 0 | 2 | 14 | 0 | 10 | 18 | 16 | 13 | 8 | 12 |
| SUB02 | 4 | 9 | 0 | 1 | 14 | 0 | 0 | 2 | 3 | 0 | 0 |
| SUB03 | 6 | 0 | 14 | 2 | 0 | 11 | 2 | 4 | 5 | 4 | 12 |
| SUB04 | 11 | 4 | 1 | 5 | 5 | 7 | 9 | 4 | 9 | 9 | 7 |
| SUB05 | 5 | 2 | 0 | 3 | 0 | 8 | 0 | 5 | 9 | 1 | 6 |
| SUB06 | 9 | 5 | 0 | 11 | 6 | 0 | 0 | 13 | 3 | 0 | 0 |
| SUB07 | 4 | 0 | 31 | 1 | 12 | 0 | 0 | 1 | 5 | 6 | 0 |
| SUB08 | 4 | 1 | 0 | 0 | 3 | 5 | 0 | 10 | 17 | 3 | 4 |
| SUB09 | 5 | 0 | 0 | 22 | 0 | 0 | 0 | 9 | 9 | 0 | 0 |
| SUB10 | 4 | 3 | 2 | 3 | 0 | 8 | 2 | 0 | 6 | 9 | 6 |
| SUB11 | 1 | 0 | 0 | 8 | 0 | 0 | 0 | 8 | 23 | 0 | 0 |
| SUB12 | 11 | 5 | 6 | 0 | 3 | 4 | 0 | 2 | 6 | 0 | 0 |
| SUB13 | 5 | 0 | 12 | 0 | 0 | 6 | 6 | 3 | 0 | 11 | 9 |
| SUB14 | 6 | 1 | 7 | 4 | 6 | 2 | 7 | 5 | 16 | 4 | 1 |
| SUB15 | 7 | 0 | 0 | 8 | 0 | 21 | 12 | 7 | 12 | 7 | 16 |
| SUB16 | 1 | 0 | 0 | 13 | 1 | 10 | 0 | 8 | 22 | 2 | 8 |
| SUB17 | 4 | 1 | 10 | 0 | 0 | 5 | 3 | 1 | 2 | 2 | 0 |
| SUB18 | 9 | 0 | 0 | 9 | 5 | 18 | 0 | 2 | 7 | 9 | 4 |
| SUB19 | 10 | 1 | 3 | 3 | 10 | 0 | 0 | 7 | 13 | 0 | 0 |
| SUB20 | 4 | 9 | 0 | 0 | 7 | 0 | 0 | 0 | 6 | 0 | 0 |
| SUB21 | 3 | 3 | 0 | 0 | 8 | 0 | 0 | 0 | 1 | 0 | 0 |
| SUB22 | 3 | 6 | 0 | 8 | 0 | 3 | 0 | 6 | 19 | 2 | 8 |
| SUB23 | 5 | 0 | 20 | 0 | 11 | 8 | 0 | 0 | 9 | 2 | 0 |
| SUB24 | 4 | 10 | 8 | 2 | 9 | 11 | 8 | 3 | 5 | 4 | 7 |

**Extended Data Table 2. Continued**
| Subject # | HPC | LFPC | CenG | FuG | IFG | IPL | IPS | ITG | MTG | PCC | Pcun |
| --- | --- | --- | --- | --- | --- | --- | --- | --- | --- | --- | --- |
| SUB25 | 1 | 0 | 6 | 8 | 8 | 2 | 0 | 16 | 6 | 0 | 0 |
| SUB26 | 1 | 0 | 8 | 2 | 0 | 7 | 1 | 1 | 7 | 5 | 19 |
| SUB27 | 3 | 0 | 3 | 4 | 9 | 0 | 0 | 6 | 13 | 0 | 0 |
| SUB28 | 4 | 0 | 0 | 12 | 6 | 4 | 0 | 12 | 10 | 2 | 2 |
| SUB29 | 6 | 2 | 0 | 31 | 6 | 12 | 1 | 18 | 33 | 9 | 4 |
| SUB30 | 3 | 0 | 4 | 2 | 0 | 0 | 0 | 0 | 4 | 0 | 0 |
| SUB31 | 6 | 0 | 0 | 0 | 0 | 3 | 0 | 1 | 13 | 0 | 2 |
| Total | 157 | 62 | 137 | 176 | 129 | 165 | 69 | 170 | 306 | 99 | 127 |

**Extended Data Table 2. Continued**
| Subject # | SPL | STG | VC | ACC | DLPFC | DMPFC | LOFC | PRE SMA | VMPFC |
| --- | --- | --- | --- | --- | --- | --- | --- | --- | --- |
| SUB01 | 1 | 5 | 12 | 1 | 6 | 0 | 0 | 1 | 0 |
| SUB02 | 0 | 0 | 0 | 5 | 7 | 10 | 16 | 0 | 6 |
| SUB03 | 0 | 0 | 4 | 0 | 0 | 0 | 0 | 7 | 0 |
| SUB04 | 0 | 11 | 0 | 7 | 3 | 0 | 0 | 1 | 2 |
| SUB05 | 0 | 1 | 9 | 1 | 2 | 0 | 5 | 0 | 4 |
| SUB06 | 0 | 11 | 0 | 0 | 2 | 0 | 0 | 0 | 2 |
| SUB07 | 0 | 0 | 0 | 6 | 13 | 3 | 0 | 14 | 0 |
| SUB08 | 0 | 8 | 0 | 0 | 5 | 0 | 17 | 0 | 5 |
| SUB09 | 0 | 7 | 8 | 0 | 0 | 0 | 2 | 0 | 0 |
| SUB10 | 0 | 7 | 24 | 0 | 4 | 0 | 0 | 0 | 3 |
| SUB11 | 0 | 4 | 0 | 0 | 4 | 2 | 0 | 0 | 7 |
| SUB12 | 0 | 9 | 0 | 9 | 8 | 4 | 0 | 1 | 10 |
| SUB13 | 0 | 0 | 0 | 2 | 3 | 0 | 10 | 13 | 2 |
| SUB14 | 0 | 3 | 0 | 0 | 5 | 4 | 0 | 0 | 0 |
| SUB15 | 3 | 13 | 22 | 1 | 6 | 4 | 0 | 0 | 1 |
| SUB16 | 1 | 12 | 3 | 7 | 0 | 0 | 6 | 0 | 8 |
| SUB17 | 0 | 0 | 0 | 2 | 0 | 0 | 7 | 0 | 9 |
| SUB18 | 0 | 3 | 21 | 0 | 3 | 0 | 0 | 0 | 0 |
| SUB19 | 0 | 0 | 0 | 8 | 15 | 6 | 23 | 3 | 3 |
| SUB20 | 0 | 0 | 0 | 5 | 21 | 4 | 14 | 3 | 19 |
| SUB21 | 0 | 0 | 0 | 7 | 14 | 12 | 1 | 7 | 2 |
| SUB22 | 0 | 12 | 11 | 0 | 3 | 0 | 0 | 0 | 2 |
| SUB23 | 0 | 0 | 0 | 4 | 10 | 3 | 0 | 7 | 6 |
| SUB24 | 5 | 9 | 8 | 2 | 30 | 19 | 13 | 7 | 9 |

**Extended Data Table 2. Continued**
| Subject # | SPL | STG | VC | ACC | DLPFC | DMPFC | LOFC | PRE SMA | VMPFC |
| --- | --- | --- | --- | --- | --- | --- | --- | --- | --- |
| SUB25 | 0 | 11 | 0 | 4 | 1 | 1 | 0 | 0 | 1 |
| SUB26 | 10 | 0 | 24 | 0 | 0 | 0 | 0 | 0 | 0 |
| SUB27 | 0 | 6 | 2 | 0 | 3 | 0 | 8 | 0 | 3 |
| SUB28 | 0 | 3 | 3 | 0 | 8 | 4 | 2 | 0 | 0 |
| SUB29 | 0 | 16 | 21 | 0 | 7 | 0 | 2 | 0 | 3 |
| SUB30 | 0 | 1 | 0 | 10 | 18 | 9 | 6 | 35 | 5 |
| SUB31 | 0 | 2 | 0 | 10 | 9 | 3 | 7 | 3 | 8 |
| Total | 20 | 154 | 172 | 91 | 210 | 88 | 139 | 102 | 120 |

## Notes

### Competing Interest Statement

The authors have declared no competing interest.

